# Coupled Environmental and Demographic Fluctuations Shape the Evolution of Cooperative Antimicrobial Resistance

**DOI:** 10.1101/2023.07.06.547929

**Authors:** Lluís Hernández-Navarro, Matthew Asker, Alastair M. Rucklidge, Mauro Mobilia

## Abstract

There is a pressing need to better understand how microbial populations respond to antimicrobial drugs, and to find mechanisms to possibly eradicate antimicrobial-resistant cells. The inactivation of antimicrobials by resistant microbes can often be viewed as a cooperative behavior leading to the coexistence of resistant and sensitive cells in large populations and static environments. This picture is however greatly altered by the fluctuations arising in volatile environments, in which microbial communities commonly evolve. Here, we study the eco-evolutionary dynamics of a population consisting of an antimicrobial resistant strain and microbes sensitive to antimicrobial drugs in a time-fluctuating environment, modeled by a carrying capacity randomly switching between states of abundance and scarcity. We assume that antimicrobial resistance is a shared public good when the number of resistant cells exceeds a certain threshold. Eco-evolutionary dynamics is thus characterized by demographic noise (birth and death events) coupled to environmental fluctuations which can cause population bottlenecks. By combining analytical and computational means, we determine the environmental conditions for the long-lived coexistence and fixation of both strains, and characterize a *fluctuation-driven* antimicrobial resistance eradication mechanism, where resistant microbes experience bottlenecks leading to extinction. We also discuss the possible applications of our findings to laboratory-controlled experiments.

## 1 Introduction

Environmental conditions, such as temperature, pH, or available resources, endlessly change over time and shape the fate of natural populations. For instance, microorganisms often live in volatile environments where resource abundance fluctuates between mild and harsh conditions, and regimes of feast alternate with periods of famine [SK98, MK18, HM20]. How environmental variability (EV), generally referring to changes not caused by the organisms themselves (e.g., supply of abiotic resources), affects species diversity is a subject of intense debate and research, see, e.g., [Gri73, MRS11, Fox13, Che94, Che00, ESAH19, MHR*et al*.21, VAME10, GPW10]. Demographic noise (DN) arising from randomness in birth and death events in finite populations is another source of fluctuations. DN is negligible in large populations and strong in small ones, where it can lead to species fixation, when one species takes over the population, or to extinction, and hence can permanently set the makeup of a community [Ewe04, CK09, BM07, WSWP23]. The dynamics of the population composition (evolutionary dynamics) is often coupled with that of its size (ecological dynamics) [Rou79], resulting in its *eco-evolutionary dynamics* [PGH09, KA14, WFM18].

When EV influences the size of a population, it also modulates the DN strength, leading to a coupling of DN and EV [WFM17, WFM18, WM20, TWAM20, SMM21, TWMA23]. This interdependence is potentially of great relevance to understand eco-evolutionary dynamics of microbial communities. The coupling of DN and EV can lead to population bottlenecks, where new colonies consisting of few individuals are prone to fluctuations [WGSV02, PW09, BBG07, Bro07], and plays an important role in the eco-evolutionary dynamics of antimicrobial resistance (AMR) [CPL^+^18, MB20].

The rise of AMR is a global threat responsible for millions of deaths [O’N16]. Understanding how AMR evolves and what mechanisms can possibly eradicate the resistance to antimicrobials are therefore questions of great societal relevance and major scientific challenges. A common mechanism of antimicrobial resistance involves the production by resistant cells, at a metabolic cost, of an extra or intracellular enzyme inactivating antimicrobial drugs [Dav94, Wri05, YCD^+^13]. When the number of resistant cells exceeds a certain threshold, there are enough toxin-inactivating enzymes, and the protection against antimicrobial drugs is shared with sensitive cells that can thus also resist antimicrobial drugs at no metabolic cost. However, below the resistant population threshold, only resistant microbes are protected against the toxin (enzyme availability is limited and it can only inactivate the drug in the vicinity of resistant cells). AMR can hence be viewed as a thresholded cooperative behavior where widespread antimicrobial inactivation is a form of public good. This results in the spread of resistant microbes below the threshold, while sensitive cells thrive under high enzymatic concentration (above threshold). Hence, in static environments and large populations, both sensitive and resistant strains survive antimicrobial treatment and coexist in the long run [YCD^+^13, VG14, MSL^+^15, BWB16]. In this work, we show that this picture can be greatly altered by the joint effect of demographic and environmental fluctuations, often overlooked, but ubiquitous in microbial communities that commonly evolve in volatile environments, where they can be subject to extreme and sudden changes [WGSV02, BBG07, Bro07, PW09, SPA^+^12, SLKF12, CPL^+^18].

Motivated by the problem of the evolution of AMR, here we study the coupled influence of EV and DN on the eco-evolutionary dynamics of a population of two species, one antimicrobial resistant strain and the other sensitive to antimicrobials. In our model, we assume that AMR is a cooperative behavior above a certain threshold for the number of resistant microbes, and the microbial community is subject to environmental fluctuations that can cause population bottlenecks. Here, EV involves random switches of the carrying capacity, causing the population size to fluctuate, while the antimicrobial input is kept constant. We thus study how the joint effect of EV and DN affects the fixation and coexistence properties of both strains, determining under which environmental conditions either of them prevail or if they both coexist for extended periods. This allows us to identify and fully characterize a *fluctuation-driven* antimicrobial resistance eradication mechanism, where environmental fluctuations generate transients that greatly reduce the resistant population and DN can then lead to the extinction of AMR.

In the next section, we introduce the model and discuss our methods. We present our results in section 3, where we first describe the main properties of the (*in silico*) model evolving under a fluctuating environment, and then study its properties analytically. In sections 3.1 to 3.3, we analyze the population dynamics in the large population limit, and then the model’s fixation properties in static environments. In section 3.4, we characterize the fixation and coexistence of the strains in fluctuating environments, and discuss in detail the fluctuation-driven eradication of antimicrobial resistance arising in the regime of intermediate switching. Section 4 is dedicated to the discussion of the influence of environmental variability on the strains fraction and abundance (section 4.1), and to a review of our modeling assumptions (section 4.2). Our conclusions are presented in section 5. Technical and computational details are given in the supplemental material (SM) [HNARM23].

## 2 Methods & Models

Microbial communities generally evolve in volatile environments: they are subject to suddenly changing conditions [SPA^+^12, SLKF12], and fluctuations can play an important role in their evolution [Gri73, MRS11, Fox13, Che94, Che00, CPL^+^18, ESAH19, MHR*et al*.21]. For instance, fluctuating nutrients may be responsible for population bottlenecks leading to feedback loops and cooperative behavior [WGSV02, BBG07, Bro07, PW09], while sensitivity to antimicrobials depends on cell density and its fluctuations [Bro04, YCD^+^13, VG14, MSL^+^15, BWB16]. Here, we study the eco-evolutionary dynamics of cooperative AMR by investigating how a well-mixed microbial community evolves under the continued application of a drug that hinders microbial growth when the community is subject to fluctuating environments. The evolutionary dynamics of the microbial community is modeled as a multivariate birth-and-death process [Gar02, vK92, BM07], whereas to model the fluctuating environment we assume that the population is subject to a time-varying binary carrying capacity [Ben06, HL06, TWAM20, TWMA23, HLGM16].

### 2.1 Microbial model

We consider well-mixed co-cultures composed of an antimicrobial resistant cooperative strain (denoted by *R*) and a defector type sensitive to antimicrobials (labeled *S*), under a constant input of antimicrobial drug. The population, of total size *N*, hence consists of *N*_*R*_ resistant and *N*_*S*_ sensitive microbes, with *N* = *N*_*R*_ + *N*_*S*_. Note that, since we later introduce EV as switches in the carrying capacity, the total population will fluctuate accordingly. A frequent mechanism of antimicrobial resistance relies on the production of an *enzyme hydrolyzing the antimicrobial drug* in their surroundings at all times [Dav94, Wri05, YCD^+^13]. When the number of *R* is high enough, the overall concentration of resistance enzyme in the medium suffices to inactivate the drug for the entire community: the enzyme hydrolyzes the drug and sets it below the *Minimum Inhibitory Concentration* (MIC), therefore acting as a public good and protecting *S* as well. This mechanism can hence lead to antimicrobial resistance as a cooperative behavior [Dav94, Wri05, YCD^+^13], for instance, by means of the *β*-lactamase resistance enzyme for the general *β*-lactam family of antibiotics [Bro09].

Here, we model this AMR mechanism by assuming that *R* acts as a cooperative strain when the number of *R* cells (proxy for resistance enzyme concentration) exceeds a fixed threshold *N*_*th*_, i.e., *R* cells are cooperators when *N*_*R*_ ≥*N*_*th*_, while they retain for themselves the benefit of producing the protecting enzyme when *N*_*R*_ < *N*_*th*_ [Bro04, YCD^+^13, VG14, MSL^+^15, BWB16]. The effective regulation of public good production by means of a population threshold has been found in a number of microbial systems, see, e.g., [BJ01, CB06, BBG07, SG13, VG14], and is consistent with a slower microbial growth cycle with respect to the fast time scale of enzyme production and dispersion. In this work, we study the AMR evolution as a form of cooperative behavior under demographic and environmental fluctuations. Assuming fixed-volume fluctuating environments, the threshold for AMR cooperation is here set in terms of *R* abundance (rather than its concentration), see section 4.

In our model, *R* microbes have a constant birth rate independent of the biostatic drug hindering microbial growth [HA12, APNK07, SMM17] ^1^, with fitness *f*_*R*_ = 1 −*s*, where 0 < *s* < 1 captures the extra metabolic cost of constantly generating the resistant enzyme. The birth rate of *S* depends on the public good abundance: when *N*_*R*_ < *N*_*th*_, the enzyme concentration is low (below cooperation threshold) and the antimicrobial drug is above the MIC, the *S* fitness *f*_*S*_ is thus lower than *f*_*R*_, with *f*_*S*_ = 1 −*a*, where 1 > *a* > *s* and *a* encodes growth rate reduction caused by the drug. When *N*_*R*_ ≥*N*_*th*_, the *R* abundance is above the cooperation threshold. This triggers the AMR cooperative mechanism: the drug is inactivated (below MIC), and the *S* birth rate, with *f*_*S*_ = 1, is then higher than that of *R*, see figure 1a. Denoting by *x* ≡ *N*_*R*_*/N* the fraction of *R* in the population, here *S* fitness is

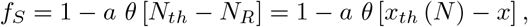

where *θ*[*z*] is the Heaviside step function, defined as *θ*[*z*] = 1 if (*z* > 0) and *θ*[*z*] = 0 otherwise, and *x*_*th*_ (*N*) ≡*N*_*th*_*/N* is the fraction of *R* at the cooperation threshold. The average population fitness is 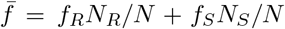. In this setting, this population evolves according to the multivariate birth-death process [Gar02, vK92, Ewe04] defined by the reactions

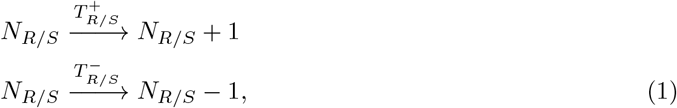

occurring with transition rates [WFM17, WFM18, TWAM20, SMM21]

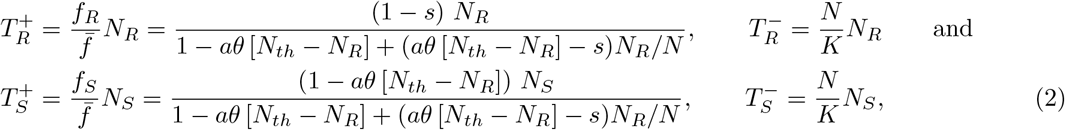

with growth limited by the logistic death rate *N/K*, where *K* is the carrying capacity, that is here assumed to be a time-fluctuating quantity, see below. Moreover, in analogy with the classical Moran model [Ewe04, BM07], we have normalized *f*_*R/S*_ by the average fitness 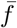.

**Figure 1:**
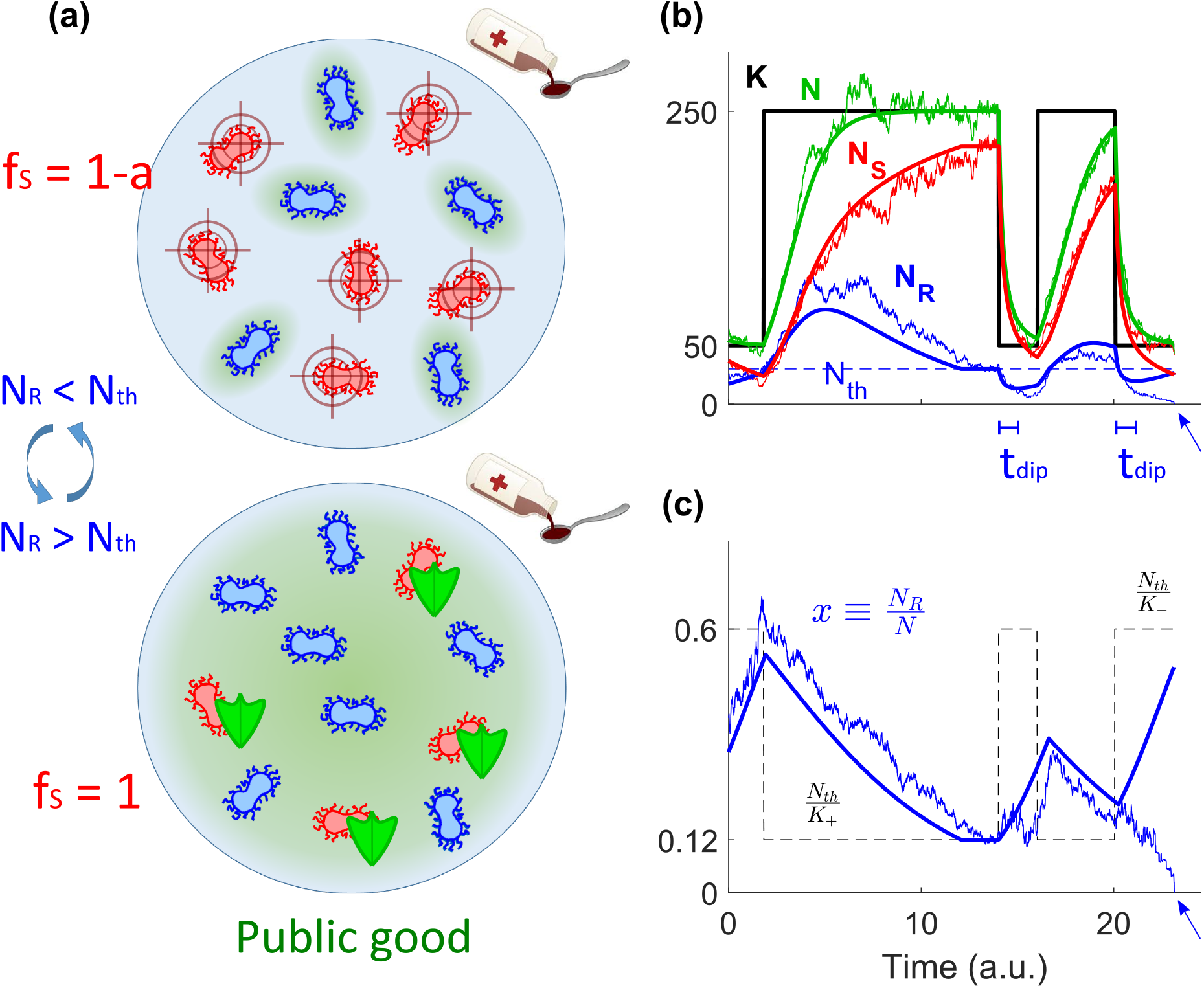
Microbial community model. **(a)** Top: When the abundance of *R* (blue microbes) is below the cooperation threshold *N*_*th*_, antimicrobial drug hinders the growth rate of *S* (red microbes) and *R* cells have a growth advantage. Bottom: AMR becomes cooperative when the number of *R* exceeds *N*_*th*_ and these generate enough *resistance enzyme* (public good in green shade) to hydrolyze the antimicrobial drug below the MIC for the whole medium, so that protection against the toxin is shared with *S* (with green shields). **(b)** Temporal eco-evolution dynamics of the microbial community for example parameters *s* = 0.2, *a* = 0.5, *K*_−_ = 50, *K*_+_ = 250, *ν* = 0.2, and *δ* = 0.6; thick black line shows the sample path of the time-switching carrying capacity *K*(*t*), with a cooperation threshold *N*_*th*_ = 30 (dashed blue line); thick solid lines depict the *N → ∞*piecewise deterministic (deterministic between two switches of *K*) process defined by equations (7) and (8) for the total microbial population (*N*, green), number of *R* (*N*_*R*_ = *Nx*, blue), and number of *S* (*N*_*S*_ = *N* (1 −*x*), red); noisy lines show an example stochastic realization of the full model under the joint effect of demographic and environmental fluctuations. In the absence of DN, *R* can experience bumps and dips (thick blue line), and *t*_*dip*_ indicates the mean time to reach the bottom of a dip from its inception; see section 3.4. In the presence of DN, fluctuations about the dip can lead to the extinction of *R* (blue arrow). **(c)** *R* fraction *x* = *N*_*R*_*/N* for the same sample path of varying environment as in (b); line styles as in panel (b); the dashed black line shows the stable *R* fraction in each environment as *K*(*t*), driven by *ξ*(*t*), switches in time.

### 2.2 Environmental Fluctuations & Master Equation

In addition to demographic fluctuations stemming from random birth and death events, see equation (1), we model environmental variability by letting the carrying capacity be a binary time-fluctuating random variable *K*(*t*) ∈{*K*_−_, *K*_+_ }, with *K*_+_ > *K*_−_. This allows us to simply model sudden extreme changes in the population size, and in particular the formation of population bottlenecks [WGSV02, PW09, BBG07, Bro07, WFM17, HLGM16].

For simplicity, we consider that *K*(*t*) is driven by the colored dichotomous Markov noise (DMN) *ξ*(*t*) = {−1, 1} that randomly switches between *K*_−_ and *K*_+_. The DMN dynamics is defined by the simple reaction [Ben06, HL06, RDL11]

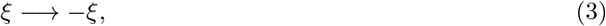

endlessly occurring at rate (1 − *δξ*)*ν*, where −1 < *δ* < 1. Here, we always consider the DMN at stationarity where *ξ* = ±1 with probability (1 ± *δ*)*/*2. The stationary DMN ensemble average is thus 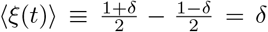 and its auto-correlation is ⟨*ξ*(*t*)*ξ*(*t*^*′*^)⟩ = (1 − *δ*^2^)*e*^−2*ν*|*t*−*t′*|^, where *ν* is both the correlation time and average switching rate. We thus consider that the binary switching carrying capacity is [WFM17, WFM17, WM20, TWAM20]

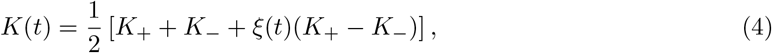

and *K*(*t*) thus switches from a state where resources are abundant (*K*_+_) to another state where they are scarce (*K*_−_) with rates *ν*_+_ ≡ *ν*(1 − *δ*) and *ν*_−_ ≡ *ν*(1 + *δ*) according to

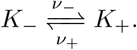

Environmental statistics can be characterized by the mean switching rate *ν* ≡ (*ν*_−_ + *ν*_+_)*/*2 and by *δ* ≡ (*ν*_−_ − *ν*_+_)*/*(*ν*_−_ + *ν*_+_) that encodes the environmental switching bias: when *δ* > 0, on average, more time is spent in the environmental state *ξ* = 1 than *ξ* = −1, and thus *K* = *K*_+_ is more likely to occur than *K* = *K*_−_ (symmetric switching arises when *δ* = 0). The time-fluctuating carrying capacity (4) modeling *environmental fluctuations* is responsible for the time-variation of the population size, and is coupled with the birth-and-death process (1) and (2).

The master equation (ME) giving the probability *P* (*N*_*R*_, *N*_*S*_, *ξ, t*) for the population to consist of *N*_*R*_ and *N*_*S*_ cells in the environmental state *ξ* at time *t* is [Gar02]:

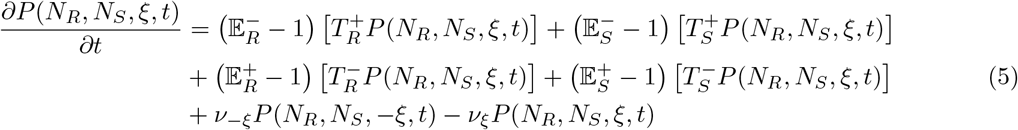

where 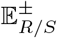 are shift operators such that 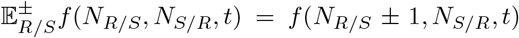, and the probabilities are set to *P* (*N*_*R*_, *N*_*S*_, *ξ, t*) = 0 whenever *N*_*R*_ < 0 or *N*_*S*_ < 0. The last line on the right-hand-side of (5) accounts for the random environmental switching, see black line in figure 1b. Since 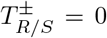 whenever *N*_*R*_ = 0 or *N*_*S*_ = 0, this indicates that there is extinction of *R* (*N*_*R*_ = 0) and fixation of *S* (*N*_*S*_ = *N*), or fixation of *R* (*N*_*R*_ = *N*) and extinction of *S* (*N*_*S*_ = 0). When one strain fixates and replaces the other, the population composition no longer changes while its size continues to fluctuate^2^. The multivariate ME (5) can be simulated exactly using standard stochastic methods (see supplemental section A [HNARM23]), and encodes the eco-evolutionary dynamics of the model whose main distinctive feature is the *coupling of the population size N and its composition x* = *N*_*R*_*/N, with DN coupled to EV*, see (2) and below.

## 3 Results

In this section we analyze how the coupled demographic and environmental fluctuations shape the evolution of the fraction of *R* in cooperative AMR [CPL^+^18]. Our main goals are to establish the conditions under which EV and DN facilitate the eradication of *R*, and reduce the size of the remaining pathogenic microbial population (see also section 4).

### 3.1 Eco-evolutionary dynamics *in silico*

The eco-evolutionary long-lived behavior of a microbial community is chiefly captured by: (I) the expected duration of the strains coexistence (Mean Coexistence Time, MCT, that here coincides with the unconditional mean fixation time [Ewe04, AS06]; see section B.2 in [HNARM23]); and (II) by the fixation (or extinction) probability of each strain, i.e., the chance that a single strain eventually takes over the entire population (or that the strain is fully replaced by others). These properties have been extensively studied in populations of constant total size, e.g., in terms of the Moran process [Mor62, Gar02, Ewe04, AS06, BM07, TH09, PRS22], but are far less known in communities of fluctuating size when DN is coupled to EV. To gain some insight into the behavior of microbial co-cultures under coupled eco-evolutionary dynamics defined by equation (5), we compute *in silico* the *R* fixation probability, denoted by *ϕ*, and the strains coexistence probability, labeled by *P*_coex_, when the external conditions fluctuate between harsh (*K*_−_ = 120, scarce resources) and mild (*K*_+_ = 1000, abundant resources). Here, *P*_coex_ is defined as the probability that both strains still coexist for a time exceeding twice the average stationary population size *t* > 2⟨*N* ⟩^3^. In our simulations, we consider a wide range of the switching rate *ν* and bias *δ*, with ∼ 10^3^ − 10^4^ realizations for each dynamic environment, and different values of the cooperation thresholds, with *N*_*th*_ ∼ 100. In our simulations, we respectively use *s* ∼ 0.1 − 0.2 and *a* ∼ 0.25 − 0.5 as plausible values for the resistance metabolic cost and the impact of the drug on *S* [vdHSS^+^11]. Our choice of *K*_*±*_ ensures that the dynamics is not dominated mainly by DN or EV, but by the interplay of DN and EV, and the values of the cooperation threshold *N*_*th*_ < *K*_−_ guarantee that the fixation of either strain or their coexistence are all scenarios arising with finite probabilities in our simulations, see below and supplemental section A [HNARM23]. Note that, as discussed in section 4.2 and supplemental section D.3, the behavior reported here can also be observed in big, realistic populations of *N* > 10^6^.

Figure 2a-c shows the *in silico ν* −*δ* phase diagrams corresponding to the various fixation and coexistence scenarios arising for different cooperation thresholds. For small thresholds relative to EV (*N*_*th*_ ≲ 10*K*_+_*/K*_−_, see supplemental section D.3 [HNARM23]), *S* displays a high fixation probability (red region) at intermediate *ν* and non-extreme *δ*, where *R* is most likely to be eradicated. Under high/low values of *ν* (when *δ* is not too low), the red region in figure 2a-c is surrounded by dark areas where the long-lived coexistence of the strains is most likely. When the threshold *N*_*th*_ is closer to *K*_−_, *R* is most likely to prevail in the blue region of figure 2b-c, where the environment is predominantly in the harsh state (*δ* < 0). As *N*_*th*_ increases, the blue region expands and gradually replaces the red and black areas: the fixation of *R* is likely to occur in most of the *ν* −*δ* diagram. In addition to the population makeup, the average population size is a decreasing function of *ν* at fixed *δ*, and increasing with *δ* at fixed *ν*; see figure 4d-e and section 4.1, and [WFM17, WFM18, TWAM20, SMM21, TWMA23].

**Figure 2:**
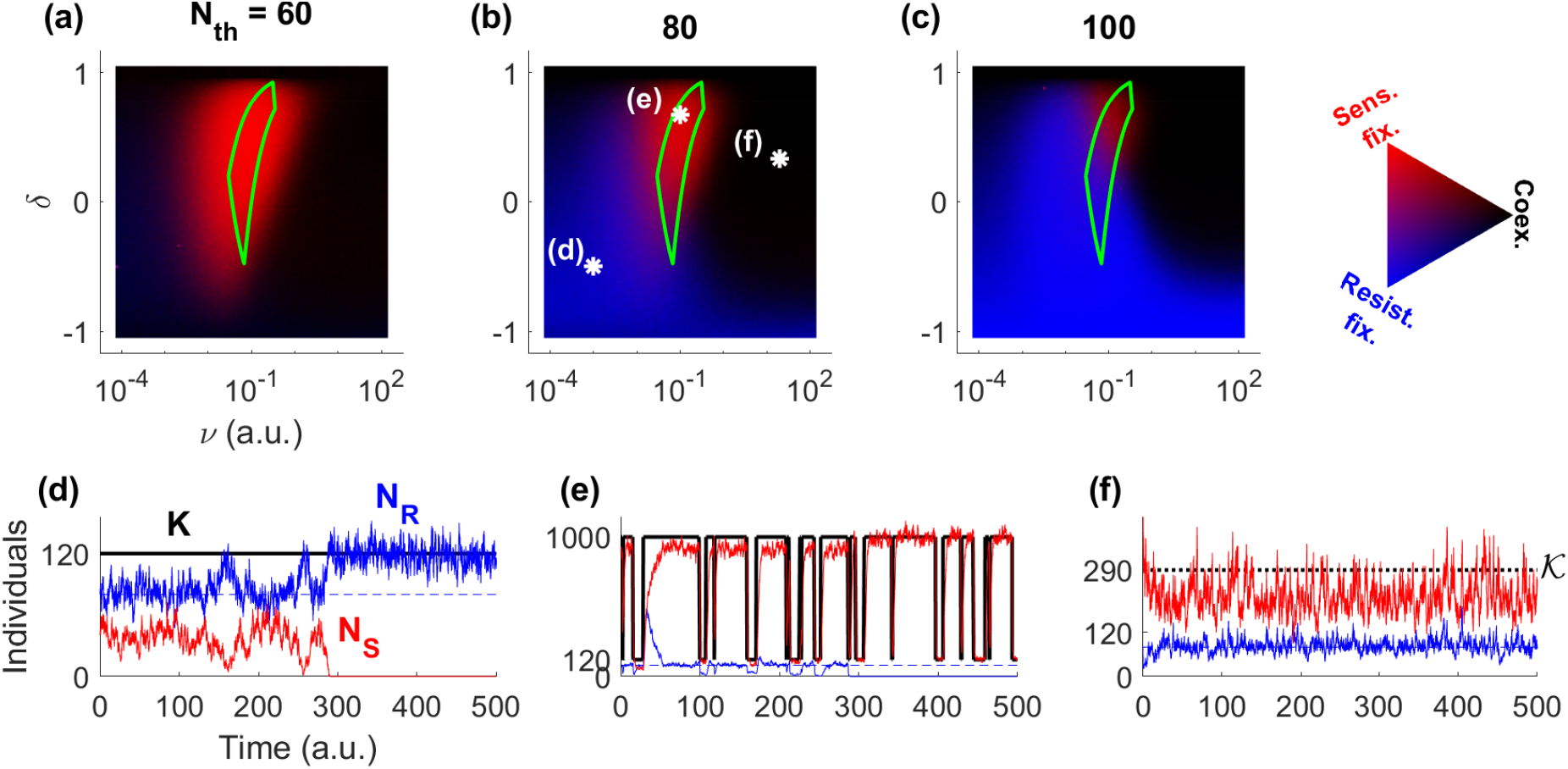
Eco-evolutionary dynamics in the phase diagram of the joint fixation and coexistence probability. **(a-c)** Fixation and coexistence joint probability *in silico* at a given environmental bias *δ* and mean switching frequency *ν* for *s* = 0.1, *a* = 0.25, *K*_−_ = 120, and *K*_+_ = 1000 at resistant cooperation thresholds *N*_*th*_ = 60, 80, and 100; see the discussions in sections 3.3, 4.2, and supplemental section D.3 [HNARM23] for the behavior at much larger populations and thresholds. Stronger blue (red) depicts a higher fixation probability of *R* (*S*). Darker color indicates higher coexistence probability, defined as the probability to not reach any fixation before *t* = 2 ⟨*N⟩*, where we take the average total population in its stationary state. The area enclosed within the green solid line indicates the optimal regime for the eradication of *R*, see section 3.4. The white asterisks in (b) depict the environmental statistics for each of the bottom panels. **(d-f)** Sample paths for the carrying capacity (*K*, black), number of *R* (*N*_*R*_, blue), number of *S* (*N*_*S*_, red), and fixed cooperation threshold *N*_*th*_ = 80 (dashed blue) for the environmental parameters (*ν, δ*) depicted by the corresponding white asterisk in (b). The high environmental switching frequency in (f) results in an effectively constant carrying capacity *K* = K; see section 3.2.

**Figure 3:**
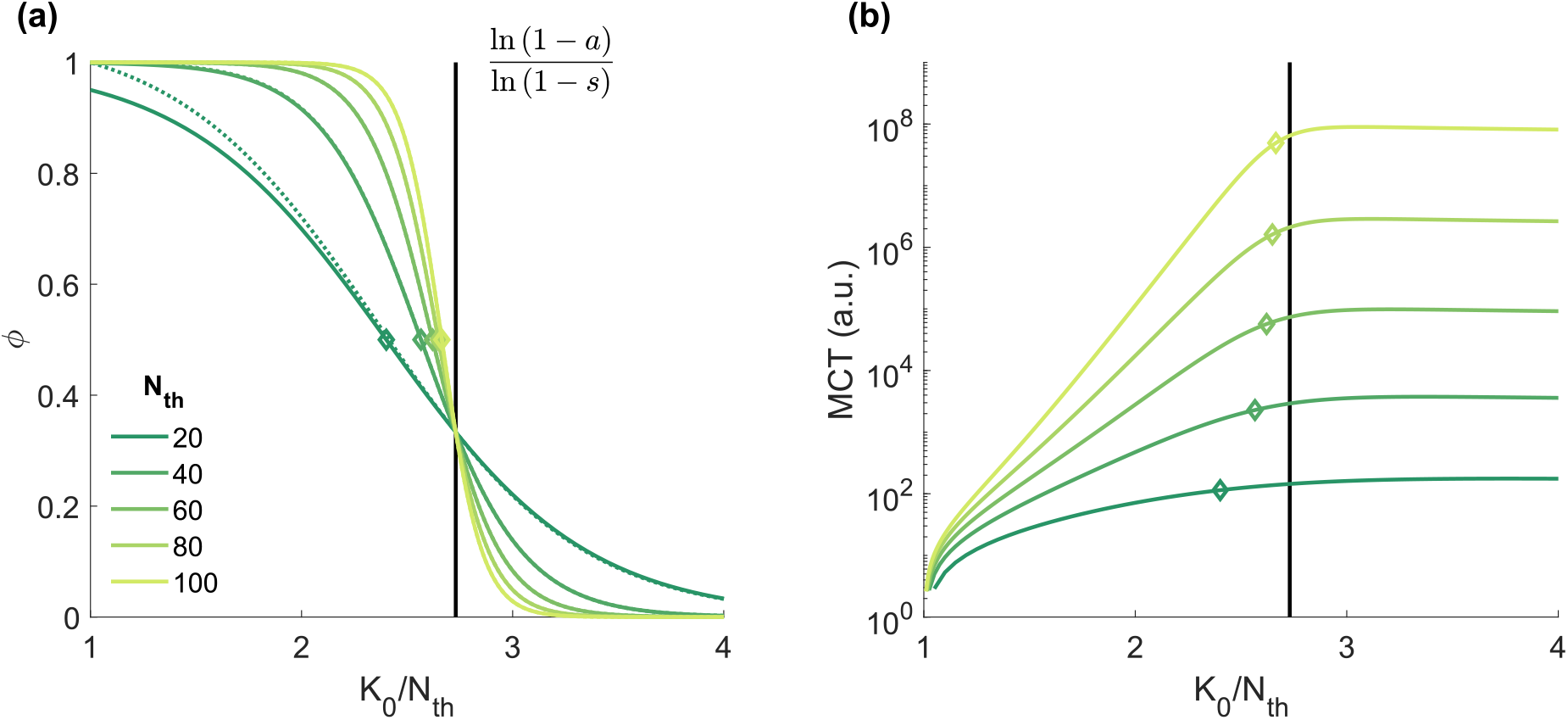
Moran theory for *R* fixation probability and Mean Coexistence Time (MCT) in static environments. **(a)** *R* fixation probability *ϕ* in terms of the total microbial population normalized by the resistant cooperation threshold *K*_0_*/N*_*th*_ for five example thresholds, from *N*_*th*_ = 20 (dark green) to 100 (yellow green); the starting microbial composition is set at the coexistence equilibrium *x*_*th*_ = *N*_*th*_*/K*_0_; solid lines show the approximated prediction of equation (9); dotted lines depict the exact Moran behavior of supplemental equation (S6) [HNARM23], only distinguishable for the smallest threshold; open diamonds illustrate the predicted 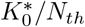 that provides fixation equiprobability for each resistant cooperation threshold, see section 3.3. **(b)** Mean Coexistence Time vs *K*_0_*/N*_*th*_ in log-linear scale; solid lines show the exact Moran MCT, computed from supplemental equation (S8); legend and symbols as in panel (a).

**Figure 4:**
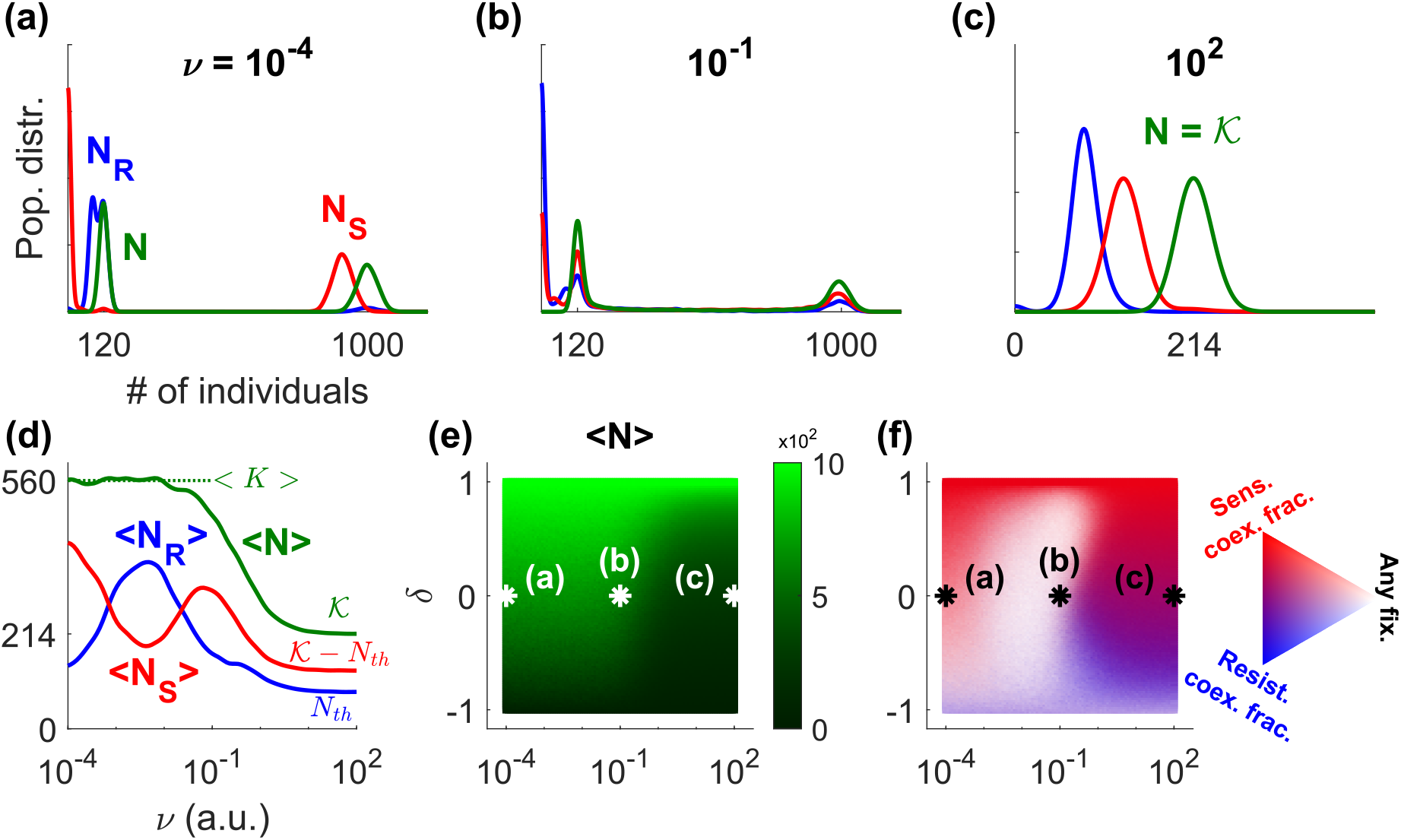
Total population, strain abundance, and coexistence composition in fluctuating environments. **(a-c)** *In silico* probability distributions of the total population (*N*, green), number of *R* (*N*_*R*_, blue), and number of *S* (*N*_*S*_, red), with parameters *s* = 0.1, *a* = 0.25, *K*_−_ = 120, *K*_+_ = 1000, and *N*_*th*_ = 80, under no environmental bias (*δ* = 0) and for mean switching rates in slow *ν* = 10^−4^ (a), intermediate *ν* = 10^−1^ (b), and fast *ν* = 10^2^ (c) conditions. Histograms are smoothed by a Gaussian filter of width *σ* = 10 in cell number. **(d)** Average overall population and strain abundances under no bias, i.e., *δ* = 0, as a function of switching rate *ν*; colors as in (a-c). Lines are smoothed by a log-scale Gaussian filter of width *σ* = 10, i.e., one frequency decade. **(e)** Average overall population size in dynamic environments. **(f)** Coexistence composition and fixation probability (of any strain) in dynamic environments. Stronger blue (red) depicts a higher coexistence fraction of *R* (*S*). Lighter color indicates lower coexistence probability, defined as the probability for no fixation event to occur before *t* = 2⟨*N* ⟩. The white and black asterisks in (e-f) depict the environmental statistics for each of the top panels. All panels are computed at quasi-stationarity reached after a time *t* = 2⟨*K*⟩, ensuring that *N* reaches its (quasi-)stationary state, where ⟨*N* ⟩ ≤ ⟨*K*⟩; see text.

In what follows, we analyze the different phases of figure 2a-c, focusing particularly on the characterization of the red area, and also determine how *N*_*R*_ varies with the environmental parameters in the different phases. This allows us to determine *the most favorable environmental conditions for the eradication of R and for the reduction of the population of pathogenic cells*, which are issues of great biological and practical relevance.

### 3.2 Sample paths

To gain an intuitive understanding of the model’s eco-evolutionary dynamics, it is useful to discuss the sample paths of figures 1b-c and 2d-f in terms of the population size *N* and the *R* fraction *x* = *N*_*R*_*/N*.

It is instructive to first consider the case of very large population arising with a constant and large carrying capacity *K*(*t*) = *K*_0_ ≫1. In this setting, corresponding to a static environment, we ignore all forms of fluctuations and the system evolves according to the mean-field (deterministic) differential equations

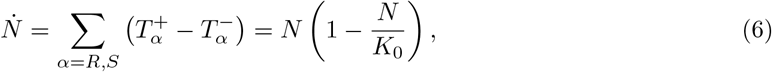

and

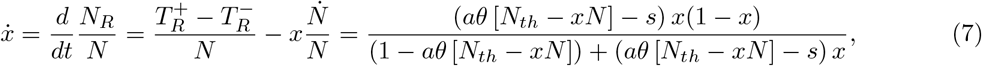

where the dot indicates the time derivative. It is clear from equation (7) that the dynamics of the population composition, given by *x*, is coupled to that of its size *N*. According to the logistic equation (6), the population size reaches *N* = *K*_0_ on a time scale *t* ∼ 1 independently of *x*, while the population composition is characterized by a stable equilibrium *x* = *x*_*th*_ ≡ *N*_*th*_*/N* = *N*_*th*_*/K*_0_ reached on a time scale of *t ∼*1*/s* or ∼1*/*(*a − s*) from *x* > *N*_*th*_*/N* or < *N*_*th*_*/N*, respectively. When *s* < *a ≪*1, there is a timescale separation, with *N* relaxing to its equilibrium much faster than *x*. We note that the coexistence equilibrium in terms of *R* and *S* is 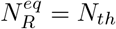 and 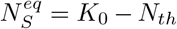 Clearly, this suggests that *S* would unavoidably be wiped out if *N*_*th*_ was greater than the carrying capacity, and hence we always consider that the latter exceeds the cooperation threshold (*K* > *N*_*th*_).

When the population is large enough for demographic fluctuations to be negligible 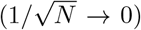 and the sole source of randomness stems from the time-fluctuating environment (random switches of the carrying capacity), the dynamics becomes a so-called piecewise deterministic Markov process (PDMP) [Dav84]. Between each environmental switch, the dynamics is deterministic and given by equations (6), with *K*_0_ replaced by *K*_*±*_ in the environmental state *ξ* = ±1, and (7). Here, the PDMP is thus defined by

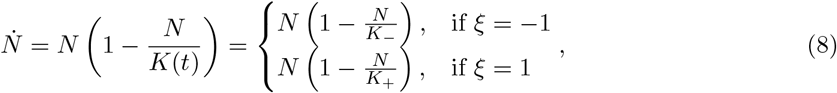

where the fluctuating carrying capacity *K*(*t*) is given by equation (4), coupled to (7). Sample paths of this PDMP are shown as solid lines in figures 1b-c and 2d-f. These realizations illustrate that *N* (*t*) tracks the switching carrying capacity *K*(*t*) independently of *x*, while *x*(*t*) evolves towards the coexistence equilibrium at the cooperation threshold *x*_*th*_(*t*) = *N*_*th*_*/N* (*t*), which changes in time as *N* varies. Hence, *x* increases when *N*_*R*_ < *N*_*th*_, and it decreases when *N*_*R*_ > *N*_*th*_. For extremely high environmental switching rate *ν → ∞*, the microbial community experiences a large number of switches, between any update of the population make-up. In this case, *N* is not able to track *K*(*t*), but experiences an effectively constant carrying capacity *K* = 𝒦 obtained by self-averaging the environmental noise (see [WFM17, WFM18, WM20, SMM21, TWMA23]):

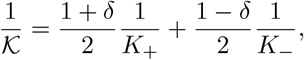

leading to 𝒦 = 2*K*_+_*K*_−_*/*[(1 − *δ*)*K*_+_ + (1 + *δ*)*K*_−_]. Hence, when *ν* → ∞, the community size is approximately *N* ≈ 𝒦 and, provided that *δ* is not too close to −1, long-lived coexistence of both strains is likely (with abundances *N*_*R*_ ≈ *N*_*th*_ and *N*_*S*_ ≈ 𝒦 − *N*_*th*_), as shown in figure 2f.

### 3.3 Fixation and coexistence in static environments (*δ* = ±1)

When EV causes a population bottleneck, DN about the coexistence equilibrium may cause the extinction of one strain and the fixation of the other (see figures 1b-c and 2d-e). To elucidate the fate of microbial communities under fluctuating environments, it is therefore necessary to first understand how a small community is able to fixate, or avoid extinction, in a static environment, when it is subject to a constant carrying capacity *K*_0_, with 1 ≪ *K*_0_ ∼ *K*_−_ ≪ *K*_+_. This condition ensures both fixation of one strain or long-lived coexistence are possible, i.e., demographic fluctuations, of order 𝒪 (1*/ K*_0_), matter but do not govern the dynamics.

Since the community composition tends to the coexistence equilibrium *x* → *x*_*th*_, see equation (7), the faster *N* dynamics reaches its steady state *N* → *K*_0_ before any fixation/extinction events occur, see equation (6). Therefore, we assume a fixed *N* = *K*_0_. The evolutionary dynamics is thus modeled by the analytically tractable Moran process [Mor62, Ewe04, BM07, CMF11, WFM17, WFM18], where the population composition evolves stochastically by balancing each birth/death of *R* by the simultaneous death/birth of a *S*, according to the reactions

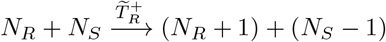

and

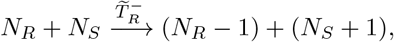

with the effective transition rates 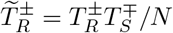 obtained from (2) [WFM17, WFM18].

Due to DN, the *R* fraction fluctuates around *x*_*th*_ until the eventual extinction of a strain. Therefore, from the classic Moran results (equation (S6) in [HNARM23]), we can derive a simplified, approximated expression for the *R* fixation probability by setting any initial composition directly at coexistence *x*_0_ = *x*_*th*_, which yields

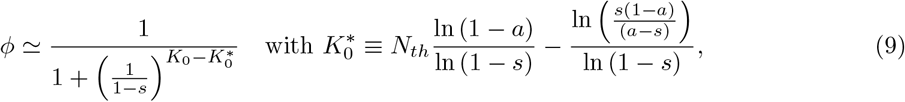

where we now assumed 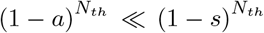 and 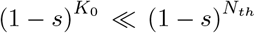, which is in line with our choices 0 < *s* < *a* < 1 and *N*_*th*_ < *K*_0_. Here 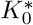 is the microbial population size giving the same fixation probability 1*/*2 to *R* and *S*. In our examples, *s* = 0.1 and *a* = 0.25 (see section 3.1), and fixation equiprobability is reached at 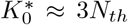, where the *R* and *S* abundance in the long-lived coexistence equilibrium are respectively 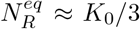and 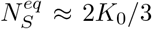. Figure 3a shows the excellent agreement between the approximation (9) (solid lines) and the exact *R* fixation probability of the underlying Moran process of supplemental equation (S6) (dotted lines), for different cooperation thresholds (*N*_*th*_ = 20 −100)^4^.

Equation (9) and figure 3a show that the final fate of cooperative AMR for typically large *in vitro* populations (e.g., *N* ≳ 10^6^ in [SG13]) in static environments is unambiguously determined by the carrying-capacity-to-threshold ratio. Namely, for fixed threshold *N*_*th*_, and up to order 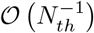, *R* is likely to fixate when *K*_0_*/N*_*th*_ < ln (1 − *a*)*/* ln (1 − *s*). On the other hand, *S* is likely to fixate when *K*_0_*/N*_*th*_ > ln (1 −*a*)*/* ln (1 −*s*). Besides the fixation of *R* or *S*, a third possible outcome is the long-lived coexistence of both strains when any fixation takes a very long time (that may be practically unobservable). The exact Moran MCT, see supplemental equation (S8) [HNARM23], shows indeed that the expected duration of coexistence increases exponentially with *N*_*th*_ and *K*_0_*/N*_*th*_, but only in the *K*_0_*/N*_*th*_ < ln (1 −*a*)*/* ln (1 −*s*) regime for the latter case; see figure 3b.

To combine the above results and understand what is the fate of the microbial community in a static environment, we remember that the mean field behavior tends to *N*_*R*_ = *N*_*th*_ and *N*_*S*_ = *K*_0_ −*N*_*th*_. When the total population is larger but close to the resistant cooperation threshold 1 ≲ *K*_0_*/N*_*th*_ < ln (1 − *a*)*/* ln (1 − *s*), *N*_*S*_ tends to be very low, *S* is thus prone to extinction and *R* to fixate. As we increase *K*_0_, *S* is less likely to go extinct and coexistence is more probable, as indicated by the exponential increase with *K*_0_*/N*_*th*_ of the MCT; see figure 3b. At populations above 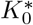 (open diamonds in figure 3), *R* has a higher extinction probability than *S* but, for cooperation thresholds *N*_*th*_ *∼*50 or higher, the mean time needed for *R* to become extinct is larger than our threshold for coexistence *t* > 2 ⟨*N⟩* = 2*K*_0_ (and generally too large to be computed in stochastic simulations; see figure 3b). We therefore conclude that, in static environments, *R* fixates when 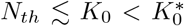, while there is long coexistence of both strains when 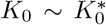 or larger. For all examples in figure 2a-c we have 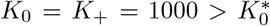 when *δ* = 1 and 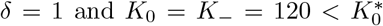 when *δ* = −1. This explains the dark areas (coexistence) in panels b-c where *δ* → 1, and the blue regions (*R* fixation) where *δ* → −1. In figure 3a, we observe dark regions (coexistence) for both *δ* = ±1 as the MCT is always larger than the coexistence threshold (*t* > 2*K*_0_). Therefore, it appears that in static environments AMR always dominates or, at least, survives for extended periods.

### 3.4 Eco-evolutionary fixation and coexistence

Under low and high environmental switching rates, the community behaves as in static total populations of size *K*_*±*_ and 𝒦, respectively; see supplemental sections D.1-D.2. Richer and novel dynamical behavior arises at intermediate switching rate (in figure 2a-c, see red areas around *ν* = 10^−2^ − 10^0^), when there are several environmental switches prior to fixation, and the quantities *ϕ* and *P*_coex_ cannot be simply expressed in terms of their counterparts in a population of constant effective size. This switching regime is characterized by the full interplay of the ecological and evolutionary dynamics: as shown in figures 1b and 2e, environmental switches can thus lead to transient “bumps” and “dips” in *N*_*R*_ (after the carrying capacity increases *K*_−_ → *K*_+_ or decreases *K*_+_ → *K*_−_, respectively). The transient *N*_*R*_ dips, together with demographic fluctuations caused by the population bottleneck (*K*_+_ →*K*_−_), can thus lead to the rapid eradication of *R* with the fixation of *S* (red areas in figure 2a-c). Each dip has a small but non-negligible probability to eradicate *R* and hence reduces the expected coexistence time.

Here, we are interested in characterizing the transient *N*_*R*_ dips as the main fluctuation-driven mechanism leading to the possible eradication of *R*. To study their properties, it is useful to consider the PDMP description of the transient *R* behavior in large populations:

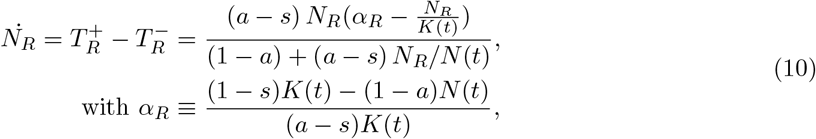

where *K*(*t*) and *N* (*t*) are respectively given by equations (4) and (8), and we assume *N*_*R*_ < *N*_*th*_. We note that, after a switch from the mild to harsh environment (*K*_+_ → *K*_−_) in the absence of DN, *R* always survives the ensuing transient dip, and *N*_*R*_ rises towards the coexistence equilibrium *N*_*R*_ = *N*_*th*_; see thick solid line in figure 1b. However, when *K*_−_ ≪ *K*_+_ and the microbial community experiences a population bottleneck, a transient dip to a small value of *N*_*R*_ can form. When this occurs, *R* is prone to extinction caused by non-negligible demographic fluctuations (stronger when *N*_*R*_ is small).

To characterize the region of the *ν*−*δ* phase diagram where transient *N*_*R*_ dips cause eradication of *R*, we need to estimate *t*_*dip*_, defined as the time from the onset of the dip to when *N*_*R*_ reaches its minimal value according to equation (10), see figure 1b. To determine *t*_*dip*_ from (10) we require 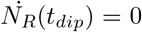 which, assuming *K*_+_ ≫ *K*_−_ ≫ 1, yields *α*_*R*_ = *N*_*R*_(*t*_*dip*_)*/K*_−_ ≈ 0, or *N* ≈ *K*_−_(1 − *s*)*/*(1 − *a*). From the solution of equation (6) with the initial condition *N* (*t* = 0) ≈ *K*_+_, we find:

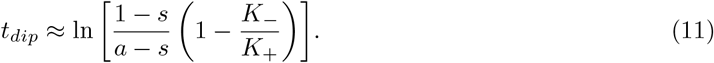

Ignoring DN, we can thus estimate the *R* population at the bottom of the transient dip 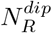, reached at *t* = *t*_*dip*_ (see supplemental section D.3 [HNARM23]):

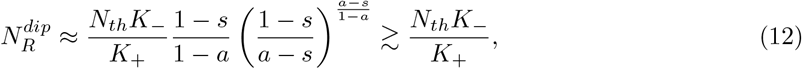

where we assumed that *R* started from *N*_*R*_(*t* = 0) = *N*_*th*_. Demographic fluctuations at the bottom of a dip are of the order 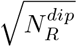. For DN to possibly drive *R* to extinction, and the fluctuation-driven eradication scenario to hold, it is necessary that 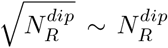, which requires 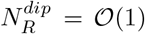, i.e. 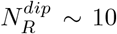 or lower. This condition is certainly satisfied when *K*_−_ and *N*_*th*_ are of comparable size (with *K*_−_ > *N*_*th*_), and each of order 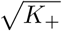, which can also hold for realistically large populations of *N* > 10^6^, see section 4.2 and section D.3 [HNARM23].

Under these sufficient requirements, the optimal environmental conditions to rapidly eradicate *R* in large but fluctuating populations can be estimated from equations (6)-(7), (10), and (11). First, in the mild environment (*K* = *K*_+_), *R* needs to be able to evolve to the coexistence equilibrium *N*_*R*_ = *N*_*th*_, requiring an average time of 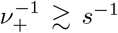. Second, after the switch from mild to harsh environment (*K*_+_ →*K*_−_), *R* needs to reach the bottom of the transient dip and experience demographic fluctuations, which imposes 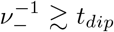. Third, if *R* survives the dip, the environment should go back to the mild *ξ* = 1 state to rule out the extinction of *S* when *ξ* = −1. For this, we require 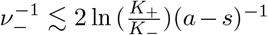, where the right-hand-side doubles the expected time to reach *N*_*R*_ = *N*_*th*_, ensuring that the dip is not interrupted by a switch; see figure 1b-c. As a fourth condition, we demand that this cycle should be as fast as possible to maximize the number of transient dips (while still allowing the population to evolve back to *N*_*R*_ = *N*_*th*_ after a bump), yielding 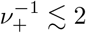 ln 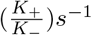, which is twice the average time needed to return to equilibrium. Using the environmental parameters *ν* and *δ*, the above lead to

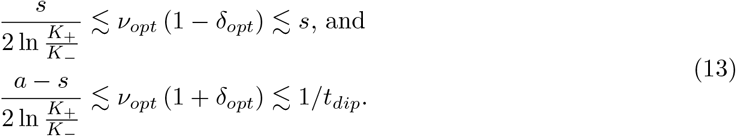

The green contour lines in figure 2a-c enclose the predicted optimal region for the fast eradication of *R* under *s* = 0.1, *a* = 0.25, *K*_−_ = 120 and *K*_−_ = 1000, and fall in the red areas observed *in silico*. The borders of these regions depend on *N*_*th*_. This stems from the dependence of *ϕ* and MCT on *N*_*th*_ (see figure 3b) and the criterion for long-lived coexistence (*t* > 2⟨*N* ⟩). The conservative prediction (13) ignores any dependence on *N*_*th*_.

## 4 Discussion

The results of the previous section characterise the long-term microbial population makeup under random switches between mild (*K* = *K*_+_) and harsh (*K* = *K*_−_) environmental conditions, for a broad range of the exogenous parameters *ν* and *δ*. Another important aspect of the time evolution of microbial population concerns the nontrivial impact of the environmental variability on the fraction and abundance of *R* and *S* in the different regimes, and especially in their phase of coexistence. It is also important to review to what extent our modeling assumptions are amenable to experimental probes.

### 4.1 Impact of environmental variability on the strains fraction and abundance

It was recently shown that in the fluctuating environment considered here, the average size of the microbial community ⟨*N* ⟩ is a decreasing function of the random switching rate *ν* (with *δ* kept fixed), that ⟨*N⟩* decreases with lower *δ* (keeping *ν* fixed), and that ⟨*N⟩ →K*_*±*_ as *δ → ±*1 [WFM17, WFM18, TWAM20, SMM21]; see figure 4d-e. As a consequence, in the blue and red areas of the phase diagrams of figure 2, where only one strain survives (figure 2a-c), the surviving pathogenic population can be reduced by increasing the environmental switching frequency *ν* and/or the time spent in harsh state (by enforcing *δ* → −1). Moreover, since the *R* fraction *x* is directly coupled to *N* through the cooperation threshold *N*_*th*_, see equation (7), environmental variability non-trivially shapes the *R* fraction in the coexistence regime (colored areas in figure 4f).

Under low switching relative to the rate of evolutionary dynamics (*ν* ≪ *s* ∼ 10^−1^ in all figures), *R* cells starting in the mild environment (*K*_+_) are able to reach the coexistence equilibrium *N*_*R*_ = *N*_*th*_ before experiencing a switch; see figure 1c. However, if the starting environment is harsh (*K*_−_), DN can rapidly eradicate *S* and destroy coexistence (see figure 2b-c). The distributions of *N*_*R*_, *N*_*S*_ and *N* in the regime *ν* → 0 are thus approximately bimodal because they combine both mild and harsh (effectively constant) environments; with *N*_*R*_ ≈ *N*_*th*_, *N*_*S*_ ≈ *K*_+_ − *N*_*th*_, and *N* ≈ *K*_+_ for the former; and *N*_*R*_ *≈K* _−_, *N*_*S*_ *≈*0, and *N ≈K*_−_ for the latter; see figure 4a. As the switching rate is increased to *ν* ≲ *s*, fixation dominates, and the *N*_*R*_ and *N*_*S*_ bimodal distributions become approximately trimodal, that is, *N*_*R/S*_ *≈*0, *K*_−_ or *K*_+_; see figure 4b. The relative weight of the peaks at *N*_*R/S*_ *≈*0 is set by *S/R* fixation probability, which is modulated by the environmental bias, see supplemental equation (S9) in [HNARM23]. The total population distribution is still bimodal about *N* = *K*_*±*_ since its relaxation dynamics (of timescale ∼ 1, see equation (6)), is faster than the evolutionary timescale ∼ 1*/s*. Finally, when *ν* is increased further (*ν* ≫ *s*), we enter the coexistence regime characterized by an effective carrying capacity *K* = 𝒦 [WFM17, WFM18, WM20, TWAM20, SMM21, TWMA23], and all distributions become unimodal about the coexistence equilibrium *N*_*R*_ *≈N*_*th*_, *N*_*S*_ *≈𝒦*−*N*_*th*_, and *N ≈𝒦*; see figure 4c.

As a consequence, if *R* is eradicated, imposing high EV (*ν ≫s*) and harsh conditions *δ →* −1 would considerably reduce the abundance of the surviving community of pathogenic *S* cells; see figure 4d, green solid line, and figure 4e. However, if *R* survives, imposing *ν ≫*1 and *δ* < 0 would not only decrease the abundance of both strains but it would also increase the *R* fraction, and risk further AMR spreading, see figure 4f (magenta /bluish areas).

### 4.2 Review of the modeling assumptions

Since we study an idealized microbial model, it is important to review our modeling assumptions in light of realistic laboratory experimental conditions. A key assumption to consider is the effectively sharp cooperation threshold *N*_*th*_, which is based on a number of experimental observations of microbial cooperation; see [Dav94, Wri05, YCD^+^13, SG13]. Accordingly, we have assumed that EV changes chemical concentrations (e.g., nutrient density) while the volume of the microbial ecosystem is kept constant [SG13]. The cooperation threshold is then fixed at a constant number of *R* microbes *N*_*R*_ = *N*_*th*_ because, at constant volume, the resistance enzyme concentration is proportional to the number of public good producers *R*. This crucial ingredient fixes the stable number of *R* at *N*_*th*_ across fluctuating environments, and is responsible for the transient dips which are at the origin of the novel eco-evolutionary mechanism for the eradication of AMR reported here. The complementary scenario, where the cooperation threshold is set by a fixed *R* fraction *x*_*th*_ is also relevant (for a different set of microbial ecosystems), and is a topic for future research.

A second assumption to review concerns the simulation results obtained here, for systems with *K*_*±*_ *∼*10^2^ −10^3^ and *N*_*th*_ *∼*100, that we are able to computationally probe (see supplemental section A [HNARM23]) but that correspond to populations of relatively small size. In the supplemental section D.3 we provide a detailed discussion on how the rich microbial behavior and novel eco-evolutionary AMR eradication mechanism reported here can be translated to larger, more realistic, microbial communities of size of order *N* ≳ 10^6^ [SG13] to *N* ≳ 10^8^ [CPL^+^18, SWJ56, Can56, Fel76, PDH^+^07], or higher. In our discussion we argue that, as long as *N*_*th*_*K*_−_*/K*_+_ ≲ 10 and 0 < *s* < *a* ≲ 10^−1^ −10^−2^, regardless of the magnitude of *K*_*±*_ or *N*_*th*_, the transient dips studied here will drag *R* close to extinction, where demographic fluctuations are instrumental for the likely and rapid eradication of AMR. Note that, for very fast/slow fluctuating environments, where transient dips are hindered (see section 3.4), *R* and *S* populations will always coexist unless *K*_−_ −*N*_*th*_ ≲ 10.

A specificity of our study is its focus on biostatic antimicrobial drugs. However, since most antimicrobials gradually change from acting as biostatic to biocidal as their concentration in the medium grows [HA12, APNK07, SMM17], our approach is consistent with a low antimicrobial concentration scenario. Conveniently, the combined biostatic effect of the drug and the normalization of strain fitness in equation (2) [Ewe04, BM07] decouples the total population *N* from its composition *x*. If any of the above conditions would not hold, *N* would then directly depend on *x*, a case already studied for a simpler model in [WFM17, WFM18]. We also note that the values used in our examples for the extra metabolic cost to generate the resistance enzyme (*s ∼*10% to 25%), and for the impact of the antimicrobial drug on *S* growth (*a ∼*25% to 50%), while only indicative, are plausible figures.

For the sake of simplicity, we have focused on binary random switching, while many laboratory experiments are often carried out with periodically switching environments, see, e.g., [AMvO08]; and natural environmental conditions often vary continuously, in time and over a range of values, see, e.g., [NLGS21]. In this context, Taitelbaum et al. studied the theoretical differences between random and periodic binary switching, and the impact of environmental fluctuations stemming from continuous environmental noise, for a prototypical two-strain model [TWAM20, TWMA23]. These studies suggest that the essence of our findings are expected to hold for general fluctuating environments with a time-varying carrying capacity, but the extent to which other and more complex forms of EV than binary random switching may alter our results for microbial communities exhibiting cooperative AMR remains a problem to be studied.

Finally, we note that the novel eco-evolutionary mechanisms reported in this study to eradicate cooperative AMR, and to reduce the total pathogenic microbial community, or minimize the coexistence fraction of *R*, all take place at a biologically and clinically relevant range of environmental switching rates. Indeed, although our theoretical study does not set a specific timescale of microbial growth, a plausible rough estimate for a single replication cycle of a microbe could be of the order of ∼1 hour. The novel AMR eradication mechanism at *ν ∼s* then comes into play when a single environmental phase lasts, on average, 1*/s ∼*10 hours. This could be consistent with the periodic administration of a treatment that enforces microbial population bottlenecks, and is a feasible time scale for laboratory experiments. Our idealized model however assumes a homeostatic influx of antimicrobial drug in all environments. Thus, an interesting approach for future work would involve the joint application of antimicrobial drug and population bottlenecks (in the harsh environment), with no drug administered in the mild environmental state.

## 5 Conclusion

Understanding how environmental variability affects the demographic and ecological evolution of microbes is central to tackle the threat of AMR, an issue of pressing societal concern [O’N16]. Central questions in studying AMR involve how the fraction of resistant microbes changes in time, and by what mechanisms these can possibly be eradicated.

It is well established that AMR is an emergent property of microbial communities, shaped by complex interactions. In particular, certain resistant cells able to inactivate antimicrobials can, under certain conditions, protect the entire microbial community. This mechanism can hence be viewed as an AMR cooperative behavior. Moreover, microbial populations are subject to changing conditions. For instance, the size of a microbial population can vary greatly with the variation of the nutrients or toxins, and can, e.g., experience bottlenecks. As a result of evolving in volatile environments, microbial communities are prone to be shaped by fluctuations. In general, these stem from environmental variability (EV, exogenous noise) and, chiefly in small populations, from demographic noise (DN). The underlying eco-evolutionary dynamics, characterized by the coupling of DN and EV, is ubiquitous in microbial ecosystems and plays a key role to understand the AMR evolution, but is still rather poorly understood.

In this work, we have studied an idealized model of cooperative AMR where a well-mixed, microbial population consisting of sensitive and resistant cells is treated with an antimicrobial (biostatic) drug, hindering microbial growth, in a fluctuating environment. The latter is modeled by a binary switching carrying capacity that fluctuates between two values corresponding to mild and harsh conditions (high/low values, respectively). Based on a body of experimental work [YCD^+^13, VG14, MSL^+^15, BWB16, Dav94, Wri05], we assume that resistant cells produce, at a metabolic cost, an enzyme that inactivates the antimicrobial drug. Importantly, the abundance of resistant microbes is thus a proxy for the concentration of the drug-inactivating enzyme, which above a certain abundance threshold, becomes a public good by providing drug protection, at no metabolic cost, to the sensitive strain. Above the cooperative threshold, the latter hence have a fitness advantage over the resistant strain, whereas, below the threshold, the drug is responsible for a reduced fitness of the sensitive cells. In this setting, the evolution of AMR can be viewed as a public good problem in a varying environment, whose outcome is shaped by the coupling of environmental and demographic fluctuations.

We have identified three regimes characterizing the eco-evolutionary dynamics of the model, associated with the fixation of the resistant or sensitive microbes, or with the long-lived coexistence of both strains. Our analysis shows that, while AMR generally survives, and often prevails, in static environments, a very different scenario can emerge under environmental variability. In fact, we demonstrate that fluctuations between mild and harsh conditions, coupled to DN, can lead to “transient dips” in the abundance of resistant microbes, which can then be driven to extinction by demographic fluctuations. Here, we determine that this *fluctuation-driven AMR eradication mechanism* occurs when the rate of environmental change is comparable to that of the relaxation of the evolutionary dynamics (*ν ∼s*). By computational means, we show that this fluctuation-driven mechanism speeds up the eradication of resistant cells, and argue that it holds also for large microbial communities, comparable to those used in laboratory experiments (*N* > 10^6^). We have also studied how EV non-trivially affects the strain abundance in the various regimes of the model, and in particular have determined the complex long-lived distribution of the fraction of resistant cells when both strains coexist and the environment fluctuates.

In conclusion, we have shown the existence of a biophysically plausible novel mechanism, driven by the coupling of EV and DN, to eradicate resistant microbes, and have demonstrated how EV shapes the long-lived microbial population in the possible scenarios of strains coexistence or fixation. Our work thus paves the way for numerous possible applications, for instance, in microbial experiments with controlled environmental fluctuations (it is currently possible to track even individual microbes, e.g., see [BLB^+^21, MSCD^+^21]), which might shed light on new possible treatments against AMR in real-world clinical infections.

## Data accessibility

Simulation data and codes for all figures are electronically available from the University of Leeds Data Repository. DOI: https://doi.org/10.5518/1360.

## Author Contributions

**Lluís Hernández-Navarro:** Conceptualization (supporting), Methodology, Formal Analysis (lead), Software, Writing - Original Draft, Writing - Review & Editing, Visualization, Investigation, Validation. **Matthew Asker:** Formal Analysis (supporting), Software, Writing - Review & Editing (supporting), Investigation, Validation. **Alastair M. Rucklidge:** Writing - Review & Editing (supporting), Supervision (supporting), Funding acquisition (supporting). **Mauro Mobilia:** Conceptualization (lead), Methodology (lead), Formal Analysis (supporting), Writing - Original Draft, Writing - Review & Editing, Visualization, Supervision (lead), Project administration, Funding acquisition (lead).

Contributor roles taxonomy by CRediT [BAA^+^15].

## Acknowledgements

We are grateful to K. Distefano, J. Jiménez, S. Muñoz Montero, M. Pleimling, M. Swailem, and U.C. Täuber for useful discussions. This work was undertaken on ARC4, part of the High Performance Computing facilities at the University of Leeds, UK.

## Footnotes

1 Here, for simplicity, we focus on biostatic drugs that reduce growth rate of sensitive cells *S*. Biocidal drugs would increase the death rate of *S*. In fact, our choice is not particularly limiting since the effect of a same drug can be either biostatic or biocidal, depending on the concentration of cells and antimicrobial [HA12, APNK07, SMM17].

2 The model will finally settle in the absorbing state *N*_*R*_ = *N*_*S*_ = 0, which corresponds to the eventual extinction of the entire population. This occurs after a time that grows exponentially with the system size and that is unobservable when, as here, *K*(*t*) ≫1 [SDF17, WFM17, WFM17, TWAM20]. This phenomenon, irrelevant for our purposes, is not considered here.

3 The rationale is that the MCT in two-strategy evolutionary games scales linearly with ⟨*N⟩* in neutral regimes, exponentially with ⟨*N⟩* in coexistence regimes, and sublinearly with ⟨*N⟩* in regimes where a strain dominates [AS06, CRF09, AM10, AM11, AHNRM23], see also [RMF07, HMT11]. Therefore, *t* > 2 ⟨*N⟩* is a conservative proxy of coexistence since it allows us to distinguish between regimes where one of the strains dominates and fixates in a time *t ≤*2 ⟨*N⟩* from a phase of long-lived coexistence (prior to the eventual fixation of one strain, after a time practically unobservable when ⟨*N* ⟩ ≫ 1).

4 We note that, in the full model simulations at static environments, the total population *N* is not fixed but fluctuates about *K*_0_. In supplemental section C [HNARM23] we discuss the minor quantitative impact of these *N* fluctuations on the fixation probability and MCT, see supplemental figure S1.

