## Supplemental Material for "Coupled Environmental and Demographic Fluctuations Shape the Evolution of Cooperative Antimicrobial Resistance"

#### A Numerical Simulations

To study the stochastic behaviour of the microbial community *in silico* we have performed *exact* stochastic simulations of the underlying birth-death process [Gil76]. Simulations start at an initial time $t = t_0$ with an initial environment $K(t_0)$ always at stationarity (with $\langle \xi(t_0) \rangle = \delta$), initial populations $N_R(t_0) = N_{th}$ and $N_S(t_0) = K(t_0) - N_{th}$, and we take into account all the possible reactions that can take place. In the case of the full model this means: (1) the four possible birth or death reactions with rates $\{T_R^+(t_0),\, T_R^-(t_0),\, T_S^+(t_0),\, T_S^-(t_0)\}$ that depend on the variables $\{N_R(t_0),\, N_S(t_0),\, K(t_0)\}$ and the constant parameters $\{s,\, a,\, N_{th}\}$; and (2) the environmental switch with constant rate $\nu_\pm$ for the state $K(t_0) = K_\pm$. We perform efficient stochastic simulations by implementing the Next Reaction Method [GB00] with an improved formulation [And07]. Simulations are run in batches of $10^3$ realizations for each constant set of parameters $\{s,\, a,\, N_{th},\, K_+,\, K_-,\, \nu_+,\, \nu_-,\, K(t_0)\}$, but for the histograms in main figure 4a-c, where we run $10^4$ to get sufficient statistical power.

We choose the lower carrying capacity as $K_- \gg 1$ but small enough to capture the impact of DN and bottlenecks on random extinctions of microbial strains; $K_- \ll K_+$ different enough so that the environmental changes have a significant impact on the dynamics; and $K_+$ as large as possible to provide insight for more realistic microbial communities, but in a finite computational time. Note that we constrain the analysis to $N_{th} < K_-$ so that microbes tend to a coexistence equilibrium in both environments, see section 3.2 in the main text. To estimate the coexistence probability $P_{\text{coex}}$ we compute the probability that strains fixate only after $t = 2\,\langle N \rangle$ (based on previous works [CRF09], see also [RMF07, HMT11]), where the expected total population size is the time average over environmental fluctuations and depends on the statistics $\nu$ and $\delta$; see main figure 4d-e. We choose this threshold, linear with $\langle N \rangle$, as a conservative proxy to distinguish coexistence and dominance regimes in small populations. The rationale is that the expected duration of coexistence $t$ in finite two-species populations scales exponentially with the system size $N$ for the former regime, whereas $t$ scales logarithmically with $N$ when there is dominance. The linear case corresponds to the neutral regime [AS06, CRF09] and separates the regimes where one species dominate from that where there is a long coexistence of both species.

Main figures 2a-c and 4f report diagrams obtained after a time $t = 2\langle N \rangle$, where $\langle N \rangle$ is the long-time mean population size. It is useful to notice that the diagrams of these figures have been obtained computationally by letting each simulation run for a time $\widetilde{t} = 2\langle K \rangle = K_+ + K_- + \delta(K_+ - K_-)$, as $\langle K \rangle$ is also the maximum value that $\langle N \rangle$ can take (for a given fixed $\delta$) [WFM17, WFM18, TWAM20]; see figure 4d. We thus record $(N_R(\widetilde{t}), N_S(\widetilde{t}))$, where any $N_{R/S}(\widetilde{t}) = 0$ implies fixation of the non-extinct strain, and coexistence corresponds to $N_{R/S}(\widetilde{t}) \neq 0$. The histograms in figure 4a-c are computed over $10^4$ realizations each, with a Gaussian filter of width $\sigma = 10$ cells to smooth the resulting curves. To computationally obtain the long-time averages of $N_R$, $N_S$, and $N$ of main figure 4d-e, we average the triplet $(N_R(\widetilde{t}), N_S(\widetilde{t}), N(\widetilde{t}))$ over $10^3$ realizations. For panel 4d, we apply a Gaussian filter of width $\sigma = 10$, i.e., one decade in the switching frequency log-scale, to smooth the curves. We also note that we obtained the simulation data reported in supplemental figure S1 by letting $10^3$ realizations run until fixation of any strain.

### B Derivations for the Moran Process

The Moran process is the stochastic 'birth-death' process where the number of individuals of two subpopulations $R$ and $S$ evolve at a strictly fixed total number $N = N_R + N_S$ [Mor62, Ewe04, BM07, AS06, TH09, WFM17, WFM18, CMF11]. The process is fully characterized by the transition rates $\widetilde{T}_R^+(N_R, N)$ and $\widetilde{T}_R^-(N_R, N)$ that quantify the rate of birth of $R$ (simultaneously balanced by a single death of $S$, $N$ being kept constant) and the rate of death of $R$ (balanced by a single birth of $S$), respectively; see main manuscript section 3.3 [WFM17, WFM18].

#### B.1 Exact General Moran Fixation Probability

The exact fixation probability $\phi\left(N_R^0, N\right)$ that the subpopulation $R$ takes over an entire population of size $N$, starting from an initial $R$ number $N_R^0$, can be derived exactly for a general two-strain Moran model with time-independent transition rates $\widetilde{T}_R^{\pm}(N_R, N)$. The exact solution for the fixation probability in the general case is [Gar02, vK92, Ewe04, AS06, TH09]

$$\phi\left(N_R^0, N\right) = \frac{1 + \sum_{k=1}^{N_R^0-1} \prod_{i=1}^{k} \gamma(i, N)}{1 + \sum_{k=1}^{N-1} \prod_{i=1}^{k} \gamma(i, N)}, \text{ for } \gamma\left(N_R^0, N\right) \equiv \frac{\widetilde{T}_R^-\left(N_R^0, N\right)}{\widetilde{T}_R^+\left(N_R^0, N\right)} \text{ and } N_R^0 = 1, 2, ..., N, \tag{S1}$$

where the factor $\gamma(N_R, N)$ fully determines the above result.

##### B.1.1 Exact particular Fixation probability

In our specific model, the effective Moran transition rates in section 3.3 of the main manuscript are $\widetilde{T}_R^+ = T_R^+ T_S^- / N$ and $\widetilde{T}_R^- = T_R^- T_S^+ / N$, obtained from the main text equation (2) [WFM17, WFM18], which read

$$\begin{aligned}
\widetilde{T}_R^+(N_R, N) &= \frac{(1-s)\cdot N_R(N-N_R)/K}{1 - a\theta\left[N_{th} - N_R\right] + (a\theta\left[N_{th} - N_R\right] - s)N_R/N}, \text{ and} \\
\widetilde{T}_R^-(N_R, N) &= \frac{(1 - a\theta\left[N_{th} - N_R\right])\cdot(N-N_R)N_R/K}{1 - a\theta\left[N_{th} - N_R\right] + (a\theta\left[N_{th} - N_R\right] - s)N_R/N}.
\end{aligned} \tag{S2}$$

Therefore, our particular factor $\gamma(N_R, N)$ yields

$$\gamma(N_R, N) = \frac{1 - a\theta\left[N_{th} - N_R\right]}{1-s}, \tag{S3}$$

which depends piecewise on the number of $R$. Substituting $\gamma$ in the general exact solution of equation (S1) we get

$$\phi(N_R^0, N) = \begin{cases} 0 & N_R^0 = 0 \\ \frac{1+\sum_{k=1}^{N_R^0-1}\left(\frac{1-a}{1-s}\right)^k}{1+\sum_{k=1}^{N_{th}-1}\left(\frac{1-a}{1-s}\right)^k+\left(\frac{1-a}{1-s}\right)^{N_{th}-1}\sum_{k=1}^{N-N_{th}}\left(\frac{1}{1-s}\right)^k} & 1 \leq N_R^0 \leq N_{th} \\ \frac{1+\sum_{k=1}^{N_{th}-1}\left(\frac{1-a}{1-s}\right)^k+\left(\frac{1-a}{1-s}\right)^{N_{th}-1}\sum_{k=1}^{N_R^0-N_{th}}\left(\frac{1}{1-s}\right)^k}{1+\sum_{k=1}^{N_{th}-1}\left(\frac{1-a}{1-s}\right)^k+\left(\frac{1-a}{1-s}\right)^{N_{th}-1}\sum_{k=1}^{N-N_{th}}\left(\frac{1}{1-s}\right)^k} & N_{th} < N_R^0 \leq 1. \end{cases} \tag{S4}$$

Note that, in consistence with the convention taken in main manuscript's section 2.1, we set $\theta[z=0] \equiv 0$. Making use of the formula for the sum of a finite geometric progression, this becomes

$$\phi(N_R^0, N) = \begin{cases} \frac{1-\left(\frac{1-a}{1-s}\right)^{N_R^0}}{1-\left(\frac{1-a}{1-s}\right)^{N_{th}}+\frac{a-s}{s(1-a)}\left(\frac{1-a}{1-s}\right)^{N_{th}}\left[\left(\frac{1}{1-s}\right)^{N-N_{th}}-1\right]} & 0 \leq N_R^0 \leq N_{th} \\ \frac{1-\left(\frac{1-a}{1-s}\right)^{N_{th}}+\frac{a-s}{s(1-a)}\left(\frac{1-a}{1-s}\right)^{N_{th}}\left[\left(\frac{1}{1-s}\right)^{N_R^0-N_{th}}-1\right]}{1-\left(\frac{1-a}{1-s}\right)^{N_{th}}+\frac{a-s}{s(1-a)}\left(\frac{1-a}{1-s}\right)^{N_{th}}\left[\left(\frac{1}{1-s}\right)^{N-N_{th}}-1\right]} & N_{th} < N_R^0 \leq 1. \end{cases} \tag{S5}$$

Finally, the fixation probability of $R$ can be written as

$$\phi\left(N_R^0, N, N_{th}, s, a\right) = \frac{1-\left(\frac{1-a}{1-s}\right)^{\frac{\left(N_R^0+N_{th}\right)-\left|N_R^0-N_{th}\right|}{2}} + \frac{a-s}{s(1-a)}\left(\frac{1-a}{1-s}\right)^{N_{th}}\left[\left(\frac{1}{1-s}\right)^{\frac{\left(N_R^0-N_{th}\right)+\left|N_R^0-N_{th}\right|}{2}}-1\right]}{1-\left(\frac{1-a}{1-s}\right)^{N_{th}} + \frac{a-s}{s(1-a)}\left(\frac{1-a}{1-s}\right)^{N_{th}}\left[\left(\frac{1}{1-s}\right)^{N-N_{th}}-1\right]}. \tag{S6}$$

An approximate simplification of the above exact result is provided in the main manuscript section 3.3 equation (9) by setting $N_R^0 = N_{th}$, $N = K_0$, and assuming $(1-a)^{N_{th}} \ll (1-s)^{N_{th}}$ and $(1-s)^{K_0} \ll (1-s)^{N_{th}}$. Example $\phi$ values for $s = 0.1$, $a = 0.25$, and several $N_{th}$ are plotted in main figure 3a and supplemental figure S1a.

##### B.2 Exact General Mean Coexistence Time (MCT)

It is also possible to exactly compute the general mean duration of coexistence regardless of the final state (either fixation or extinction of $R$), i.e., the Mean Coexistence Time (MCT) $t\left(N_R^0, N\right)$, when the transition rates $\widetilde{T}_R^{\pm}$ are time-independent. It is worth noting that the MCT here coincides with the unconditional mean fixation time (and mean extinction), since $t\left(N_R^0, N\right)$ gives the mean time after which coexistence is lost due to the fixation of one strain and the extinction of the other.

The exact formula for the MCT reads [Gar02, vK92, Ewe04, AS06, TH09]:

$$t\left(N_R^0, N\right) = -\left[\frac{\sum_{k=1}^{N-1}\sum_{n=1}^{k}\frac{\prod_{m=n+1}^{k}\gamma(m,N)}{\widetilde{T}_R^{+}(n,N)}}{1+\sum_{k=1}^{N-1}\prod_{i=1}^{k}\gamma(i,N)}\right]\sum_{k=N_R^0}^{N-1}\prod_{i=1}^{k}\gamma(i,N) + \sum_{k=N_R^0}^{N-1}\sum_{n=1}^{k}\frac{\prod_{m=n+1}^{k}\gamma(m,N)}{\widetilde{T}_R^{+}(n,N)}, \tag{S7}$$

$$\text{for } N_R^0 = 1, 2, ..., N.$$

where $\gamma\left(N_R^0, N\right)$ and $\widetilde{T}_R^{\pm}\left(N_R^0, N\right)$ are defined as in equations (S1) and (S2).

For clarity, we split in two the sum over $k$ in the first numerator (from $k = 1$ to $N_R^0 - 1$, and from $k = N_R^0$ to $N-1$), and then rearrange the equation as

$$t\left(N_R^0, N\right) = \phi\sum_{k=N_R^0}^{N-1}\sum_{n=1}^{k}\frac{\prod_{m=n+1}^{k}\gamma(m,N)}{\widetilde{T}_R^{+}(n,N)} - [1-\phi]\sum_{k=1}^{N_R^0-1}\sum_{n=1}^{k}\frac{\prod_{m=n+1}^{k}\gamma(m,N)}{\widetilde{T}_R^{+}(n,N)}. \tag{S8}$$

where $\phi \equiv \phi\left(N_R^0, N\right)$ is the $R$ fixation probability starting from $N_R(t=0) = N_R^0$ in a population of overall size $N$, and is given by (S1).

##### B.3 Coexistence probability

To derive an expression for the coexistence probability $P_{\text{coex}}$ at a fixed total population $N$, cooperation threshold $N_{th}$, and starting with the number of $R$ cells at equilibrium $N_R^0 = N_{th}$, we first compute the exact Moran MCT $t(N_{th}, K_0)$ from equation (S8); see main figure 3b and supplemental figure S1b for $N = K_0$, dotted lines. Since the microbial community tends to a coexistence equilibrium, the fixation of a strain occurs on a slow time scale, driven by fluctuations. We thus assume that the full density of coexistence times roughly approximates an exponential distribution of mean $t(N_{th}, K_0)$, a known property of systems exhibiting metastability [AM17]. $P_{\text{coex}}$ is thus the exponential cumulative probability remaining after an elapsed time $2K_0$, i.e., $P\left(t > 2\langle N\rangle = 2K_0\right)$, as previously computed *in silico* (but for a wide range of $\nu$); see main section 3.1 and figure 2a-c.

#### C Full model in a static environment with constant carrying capacity

In this section we relax the Moran approximation of a fixed total population size, and allow $N$ to fluctuate around a constant capacity, here denoted by $K_0$. In this case, the behavior of the microbial

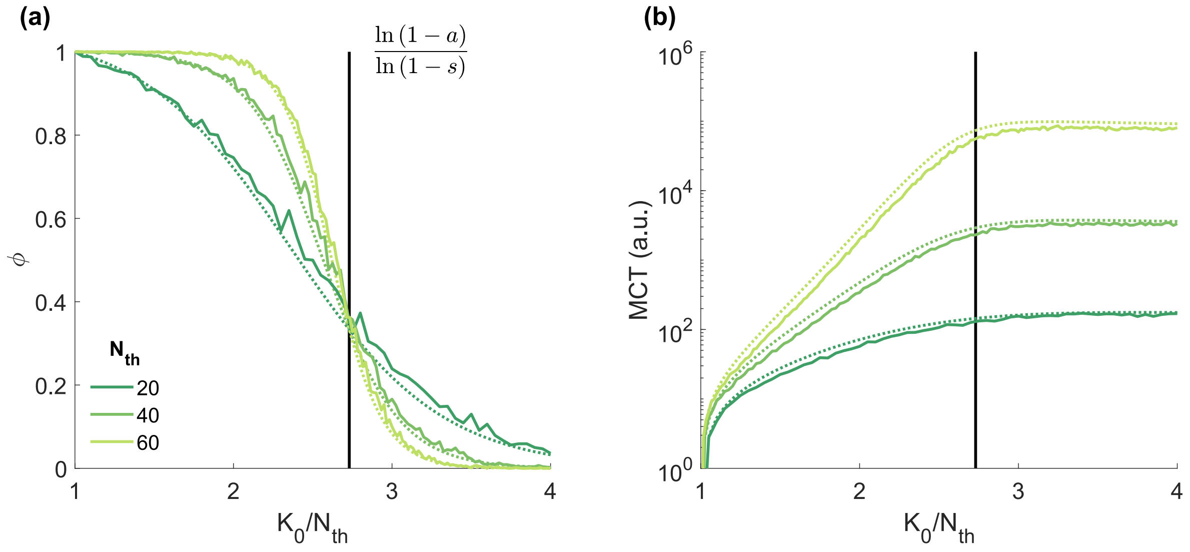


Figure S1: **Full model simulations in static environments against exact Moran theory for $R$ fixation probability and Mean Coexistence Time (MCT).** **(a)** $R$ fixation probability $\phi$ in terms of the total microbial population normalized by the resistant cooperation threshold $K_0/N_{th}$ for three example thresholds, $N_{th} = 20$ (dark green), 40 (green), and 60 (yellow green); the starting microbial composition is set at the coexistence equilibrium $x_0 = x_{th} = N_{th}/K_0$; dotted lines depict the exact Moran behavior of equation (S6), solid lines show simulation data of the full model averaged over $10^3$ realizations. **(b)** Mean Coexistence Time vs $K_0/N_{th}$ in log-linear scale; dotted lines show the exact Moran MCT, computed from equation (S8), solid lines show averaged simulation data of the full model over $10^3$ runs; legend and symbols as in panel (a).

community does not directly correspond to the Moran process, and follows a bivariate process in terms of the number $N_{R/S}$ of $R/S$ individuals (but it does not depend on $\xi$ since the environment is here static). The probability $P(N_R, N_S, t)$ that the population consists of $N_R$ and $N_S$ at time $t$, now satisfies the ME

$$\frac{\partial P(N_R, N_S, t)}{\partial t} = \left(\mathbb{E}_R^- - 1\right)\left[T_R^+ P(N_R, N_S, t)\right] + \left(\mathbb{E}_S^- - 1\right)\left[T_S^+ P(N_R, N_S, t)\right] + \left(\mathbb{E}_R^+ - 1\right)\left[T_R^- P(N_R, N_S, t)\right] + \left(\mathbb{E}_S^+ - 1\right)\left[T_S^- P(N_R, N_S, t)\right].$$

where the transition rates $T^{\pm}_{R/S}$ are given by main manuscript, equation (2), and the carrying capacity is now constant as $K \to K_0$.

For direct comparison, in figure S1 we display exact simulations of the above behavior on top of the exact Moran results of main figure 3 (dotted lines). We observe that the theoretical predictions of the Moran process quantitatively capture the results *in silico* with some minor systematic deviations. These small discrepancies arise from the fact that, below $K_0^*/N_{th} = \ln(1-a)/\ln(1-s) \approx 3$ (see main manuscript, section 3.3), the MCT depends exponentially on $K_0/N_{th}$. Therefore, it is exponentially more probable to observe faster fixation under smaller populations, i.e., when random fluctuations drive $N$ below its expected value $\langle N \rangle = K_0$.

The Moran approximation, based on assuming $N = K_0$, misses demographic fluctuations of order $\sqrt{K_0}$ about $K_0$, which results in underestimating $\phi$ and overestimating the MCT. The relative amplitude of these deviations scale with those of the standard deviations of $N/K_0$ which are of order $\mathcal{O}(\sqrt{K_0}/K_0)$ and hence become vanishingly small when $K_0 \to \infty$.

Hence, the analytical predictions from the Moran process at $N = K_0$ are relevant for static environments because they quantitatively capture both the fixation probability and MCT *in silico* within a small relative error. Moreover, this error decreases the bigger the total population and cooperation threshold as the fluctuations about fixed $K_0/N_{th}$ become negligible, which is the case for more realistic, biologically plausible microbial populations.

### D Dynamic environments: additional analytical derivations

#### D.1 Infrequent environmental switching limit $\nu \to 0$

When $\nu \to 0$, the average number of environmental switches prior to fixation of one strain (extinction of the other) is very low, and the fixation probability can be obtained by averaging its constant-$N$ counterpart over the stationary distribution of $\xi$. We can indeed assume that the community evolves subject to a static carrying capacity set by the starting environment $K(t=0) \equiv K_0$, where $K_0 = K_\pm$ with probability $(1 \pm \delta)/2$. For these infrequently switching environments, the $R$ fixation probability at an arbitrary environmental bias $\delta$ is the average

$$\phi(\nu \to 0, \delta) = \frac{(1+\delta)\phi(K_+) + (1-\delta)\phi(K_-)}{2}, \tag{S9}$$

where $\phi(K_\pm) \equiv \phi(N_{th}, K_\pm)$ is the Moran fixation probability of the exact equation (S6), or approximate main equation (9), starting at the equilibrium $N_R^0 = N_{th}$. As for the example $s = 0.1$ and $a = 0.25$ parameter values shown in main figures 2-4, this $\phi(\nu \to 0, \delta)$ $R$ fixation probability is high for $K_0 = K_- = 120$ with $\delta = -1$, and gradually (linearly) lowers as $\delta$ increases and the weighted average shifts towards the starting environment $K_0 = K_+ = 1000$ at $\delta \to +1$; see figure 3a at the limiting cases $K_\pm/N_{th}$, with $N_{th} \in [60, 100]$; and see the blue-to-black gradient when $\nu \to 0$ for increasing $\delta$ in figure 2a-c.

Similarly as discussed above for the fixation probability, the coexistence probability at infrequently switching environments averages across the two possible initial environments as

$$P_{\text{coex}}(\nu \to 0, \delta) = \frac{(1+\delta)P_{\text{coex}}(K_+) + (1-\delta)P_{\text{coex}}(K_-)}{2}, \tag{S10}$$

where $P_{\text{coex}}(K_\pm)$ is the coexistence probability in a static environment $K_\pm$, at fixed total population $N = K_\pm$, and starting $R$ population $N_R^0 = N_{th}$, as derived in the previous supplemental section B.3. The slow switching environment coexistence probability $P_{\text{coex}}(\nu \to 0, \delta)$ of the above equation (S10) is small for low $\delta \to -1$ and linearly larger for high $\delta \to +1$. This is because the most likely (initial) value of the carrying capacity varies linearly with $\delta$, from $K_0 = K_-$ when $\delta \to -1$ to $K_+$ for $\delta \to 1$; see the MCT in main figure 3b at $K_\pm/N_{th}$ compared to the corresponding coexistence duration threshold $2K_\pm$. Therefore, in dynamic environments with very infrequent switches, microbial behavior shifts from fast fixation (bright) of $R$ (blue) to very slow fixation (black) of $S$, interpreted here as long-lived coexistence (red, overshadowed by black); see section 3.3 and figure 3 at $K_\pm/N_{th}$. Note that the MCT at $K(\delta \to -1) = K_-$ is largest for the smallest threshold $N_{th} = 60$ (see figure 3b), so that the bright blue region in figure 2a is overshadowed by coexistence, in black.

#### D.2 Frequent environmental switching limit $\nu \to \infty$

When $\nu \to \infty$, the carrying capacity experiences numerous switches before fixation and extinction occurs. This results in the self-averaging of the environmental noise and the total microbial population tends to the effective carrying capacity $\mathcal{K}$ seen in section 3.2 [WFM17, WFM18, WM20, TWAM20, SMM21, TWMA23]:

$$N \to \mathcal{K}(\delta) \equiv \frac{2K_+K_-}{(1-\delta)K_+ + (1+\delta)K_-},$$

with $K(\delta \to \pm 1) = K_\pm$. Therefore, the theoretical $R$ fixation and coexistence probabilities in the high environmental switching frequency limit are effectively those of a static environment with $K_0 = \mathcal{K}$, that is

$$\phi(\nu \to \infty, \delta) = \phi(\mathcal{K}(\delta)), \text{ and } P_{\text{coex}}(\nu \to \infty, \delta) = P_{\text{coex}}(\mathcal{K}(\delta)). \tag{S11}$$

where $\phi(\mathcal{K}(\delta))$ and $P_{\text{coex}}(\mathcal{K}(\delta))$ are the static environment $\mathcal{K}$, fixed total population $N = \mathcal{K}$, starting at equilibrium $N_R^0 = N_{th}$, fixation and coexistence probabilities of the exact equation (S6) (or approximate main equation (9)) and section B.3, respectively.

Consistent with the *in silico* results of figure 2a-c with $\nu \to \infty$, this limiting behavior also introduces a blue-to-black (nonlinear) gradient for $\delta = -1 \to +1$ as the effective carrying capacity gradually shifts from $\mathcal{K} = K_-$ to $K_+$. As a result, we observe a sharp transition from fixation of $R$ (bright blue) to long-term coexistence (black), with the eventual slow fixation of $S$; see figure 3 at $\mathcal{K}(\delta)/N_{th}$.

#### D.3 Realistic population numbers $N > 10^6$

Typical microbiology laboratory experiments study total microbial populations of typical size $N \sim 10^6$ or bigger [SG13]. These studies model real-life microbial communities that are usually a few orders of magnitude larger, such as in case studies of mature or chronical clinical infections with $N \gtrsim 10^8$ [CPL+18, SWJ56, Can56, Fel76, PDH+07]. In our study, we are constrained to consider systems that are amenable to scrutiny over a wide range of environmental parameters $\{\nu, \delta\}$ in feasible computational time, and have hence restricted our *in silico* simulations to populations of size up to $N = 1000$. It is thus important to assess analytically how our main findings *in silico* translate to microbial population of more realistic size $N > 10^6$.

To this end, we first notice that, as discussed in section 4 of the main text, the environmental parameters must fulfill $1 < K_-/N_{th}$ to avoid that $S$ dives into extinction following a switch to the harsh environment. For this, the stable number of $S$, $N_S = K_- - N_{th}$ (see section 3.2 in the main text), must be high to resist demographic fluctuations, for which a reasonable estimate is $K_- - N_{th} > 10$. The second point to notice is that the model tends to two possible coexistence equilibria $0 < x = N_{th}/K_\pm < 1$ as long as $K_- > N_{th}$ (see section 3.2). Hence, the expected behavior for large populations in the two possible static environments $K_\pm$ is coexistence, which has a duration that scales exponentially with the cooperation threshold $N_{th}$; see main figure 3b. Since typically the resistant cooperation threshold is of the order $N_{th} \lesssim K_-$, the duration of microbial coexistence is thus expected to grow exponentially with $K_-$. Therefore, fixation in static environments is never observed in realistically big communities where $K_- \gtrsim 10^6$.

However, as discussed in the main section 3.4, the coupled eco-evolutionary dynamics in fluctuating environments generates significant transient $N_R$ dips when switching from mild to harsh environments ($K_+ \to K_-$) at an intermediate switching rate $\nu \sim s$. The frequency and depth of these transient dips are maximized in a certain range of environmental parameters $\nu$ and $\delta$, derived in section 3.4. The rapid eradication of $R$ in this optimal dynamic environment regime is shown in the green-enclosing areas of figure 2a-c. We now ask whether realistically big values of $\{N_{th}, K_-, K_+\}$ actually enhance the extinction of cooperative AMR in the fluctuation-driven optimal AMR eradication regime $\{\nu, \delta\}$.

In such an optimal AMR-eradication regime, the minimum possible expected number of $R$, reached in the transient dips, is given by the main equation (12). To derive this equation, on the one hand we take the low $R$ fraction limit $x \to 0$ in the main equation (7)

$$\dot{x} \approx \frac{a-s}{1-a} x,$$

and thus, assuming $x(t=0) = x_+^{eq} \equiv \frac{N_{th}}{K_+}$, where $t = 0$ is the time at the $K_+ \to K_-$ environmental switch, we obtain the microbial composition dynamics at short times

$$x(t) \approx \frac{N_{th}}{K_+} e^{\frac{a-s}{1-a} t}.$$

On the other hand, we can exactly solve the total population logistic dynamics of main equation (6) in the $K_- \gg 1$ environment after the switch, with $N(0) = K_+$, as

$$N(t) = \frac{K_+ K_- e^t}{K_+ (e^t - 1) + K_-}.$$

Taking $K_-/K_+ \ll 1$ in $t_{dip}$ given by main equation (11); evaluating $x(t = t_{dip})$ above; noting that $N(t = t_{dip}) \equiv K_-(1-s)/(1-a)$ from $\alpha_R \simeq 0$ in main equation (10), see main section 3.4; and multiplying both resulting expressions, provides the final estimate of the $R$ number $N_R^{dip}$ at the bottom of the transient dip:

$$N_R^{dip} = x(t_{dip}) N(t_{dip}) \approx \frac{N_{th} K_-}{K_+} \frac{1-s}{1-a} \left(\frac{1-s}{a-s}\right)^{\frac{a-s}{1-a}}. \quad \text{(S12)}$$

Since we focus on the case where DN eradicates $R$, for the transient-dip fluctuation-driven eradication mechanism to possibly work, we then need large demographic fluctuations (of order $\sqrt{N_R^{dip}}$) relative to $N_R^{dip}$. This typically suggests to consider $N_R^{dip} \sim 10$ or lower.

Big cooperative AMR microbial communities in ecosystems with antimicrobial drugs could present populations of, for instance, $N \approx K_+ \sim 10^{12}$ in nutrient abundance conditions. Moreover, biophysically plausible values for the remaining parameters could be $s = 0.1$, $a = 0.25$, and a resistant cooperation threshold $N_{th} = 2 \cdot 10^6$ for an example fixed environmental volume. Sudden and drastic ecological bottlenecks in events of nutrient scarcity (or additional toxins) could then kill most of the community and reduce the total population by a factor of, e.g., few in a million; where the total population would decrease from $N \sim 10^{12}$ to $N \approx K_- \sim 5 \cdot 10^6$. These realistic parameters would fulfill the condition $1 < K_-/N_{th} = 2.5$ with $K_- - N_{th} \sim 10^6 \gg 10$. Crucially, these plausible values would fulfill $N_R^{dip} \approx 1.7 \cdot 10 \lesssim 10$ in equation (S12), so that there would be a significant chance that $R$ becomes extinct during each transient $N_R$ dip in dynamic environments.

Finally we note that, as shown above, the relative magnitude of the population bottleneck $K_-/K_+$ is critical to enhance the extinction of $R$ during transient dips. Any increase in the population drop between abundance and scarcity environments, i.e., $K_-/K_+ \to 0$, boosts the eradication of AMR; whereas a reduction in the drop size, i.e., $K_-/K_+ \to 1$, hinders $R$ extinction and promotes strain coexistence. Tuning the nutrient abundance or scarcity levels in each environment modulates the relative magnitude of the bottleneck $K_-/K_+$. But, additionally, introducing an intermediate environmental step between harsh and mild regimes could reduce the population bottleneck and boost coexistence [SG13].

[BM07] RA Blythe and AJ McKane. Stochastic models of evolution in genetics, ecology and linguistics. *J. Stat. Mech.*, P07018, 2007.

[Can56] G Canetti. Dynamic aspects of the pathology and bacteriology of tuberculous lesions. *American Review of Tuberculosis and Pulmonary Diseases*, 74(2-2):13–21, 1956.

[CMF11] J Cremer, A Meilbinger, and E Frey. Evolutionary and population dynamics: a coupled approach. *Phys. Rev. E*, 84:051921, 2011.

[CPL+18] J Coates, BR Park, D Le, Şimşek E, W Chaudhry, and M Kim. Antibiotic-induced population fluctuations and stochastic clearance of bacteria. *eLife*, 7:e32976, 2018.

[CRF09] J Cremer, T Reichenbach, and E Frey. The edge of neutral evolution in social dilemmas. *New Journal of Physics*, 11(9):093029, 2009.

[Ewe04] WJ Ewens. *Mathematical Population Genetics.* Springer, New York, 2004.

[Fel76] WE Feldman. Concentrations of bacteria in cerebrospinal fluid of patients with bacterial meningitis. *The Journal of Pediatrics*, 88(4):549–552, 1976.

[Gar02] CW Gardiner. *Handbook of Stochastic Methods.* Springer, USA, 2002.

[GB00] MA Gibson and J Bruck. Efficient exact stochastic simulation of chemical systems with many species and many channels. *The Journal of Physical Chemistry A*, 104(9):1876–1889, 2000.

[Gil76] DT Gillespie. A general method for numerically simulating the stochastic time evolution of coupled chemical reactions. *J. Comput. Phys.*, 202:403, 1976.

[HMT11] Q He, M Mobilia, and UC Täuber. Coexistence in the two-dimensional May-Leonard model with random rates. *Eur. Phys. J. B*, 82:97–105, 2011.

[Mor62] PAP Moran. *The Statistical Processes of Evolutionary Theory.* Oxford, UK: Clarendon, 1962.

[PDH+07] M Palaci, R Dietze, DJ Hadad, FKC Ribeiro, RL Peres, SA Vinhas, ELN Maciel, V do Valle Dettoni, L Horter, WH Boom, et al. Cavitary disease and quantitative sputum bacillary load in cases of pulmonary tuberculosis. *Journal of Clinical Microbiology*, 45(12):4064–4066, 2007.

[RMF07] T Reichenbach, M Mobilia, and E Frey. Mobility promotes and jeopardizes biodiversity in rock–paper–scissors games. *Nature*, 448:1046–1049, 2007.

[SG13] A Sanchez and J Gore. Feedback between population and evolutionary dynamics determines the fate of social microbial populations. *PLoS Biology*, 11(4):e1001547, 2013.

[SMM21] S Shibasaki, M Mobilia, and S Mitri. Exclusion of the fittest predicts microbial community diversity in fluctuating environments. *J. R. Soc. Interface*, 18:20210613, 2021.

[SWJ56] MR Smith and WB Wood Jr. An experimental analysis of the curative action of penicillin in acute bacterial infections: Iii. the efffect of suppuration upon the antibacterial action of the drug. *The Journal of Experimental Medicine*, 103(4):509–522, 1956.

[TH09] A Traulsen and C Hauert. Stochastic evolutionary game dynamics. *Reviews of Nonlinear Dynamics and Complexity*, 2:25–61, 2009.

[TWAM20] A Taitelbaum, R West, M Assaf, and M Mobilia. Population dynamics in a changing environment: random versus periodic switching. *Phys. Rev. Lett.*, 125(4):048105, 2020.

[TWMA23] A Taitelbaum, R West, M Mobilia, and M Assaf. Evolutionary dynamics in a varying environment: Continuous versus discrete noise. *Physical Review Research*, 5(2):L022004, 2023.

[vK92] NG van Kampen. *Stochastic Processes in Physics and Chemistry.* North-Holland, Amsterdam, 1992.

[WFM17] K Wienand, E Frey, and M Mobilia. Evolution of a fluctuating population in a randomly switching environment. *Phys. Rev. Lett.*, 119(15):158301, 2017.

[WFM18] K Wienand, E Frey, and M Mobilia. Eco-evolutionary dynamics of a population with randomly switching carrying capacity. *J. R. Soc. Interface*, 15(145):20180343, 2018.

[WM20] R West and M Mobilia. Fixation properties of rock-paper-scissors games in fluctuating populations. *J. Theor. Biol.*, 491:110135, 2020.
